# Metabolic handoffs between multiple symbionts may benefit the deep-sea bathymodioline mussels

**DOI:** 10.1101/2023.02.09.527947

**Authors:** Tal Zvi-Kedem, Simina Vintila, Manuel Kleiner, Dan Tchernov, Maxim Rubin-Blum

## Abstract

Bathymodioline mussels rely on thiotrophic and methanotrophic chemosynthetic symbionts for nutrition, yet, secondary heterotrophic symbionts are often present and play an unknown role in the fitness of the organism. The bathymodioline *Idas* mussels that thrive in gas seeps and on sunken wood in the Mediterranean Sea and the Atlantic Ocean, host at least six symbiont lineages that often co-occur, including the primary, chemosynthetic methane- and sulfur-oxidizing gammaproteobacteria, and the secondary Methylophagaceae, Nitrincolaceae and Flavobacteraceae symbionts, whose physiology and metabolism are obscure. Little is known about whether and how these symbionts interact or exchange metabolites. Here we curated metagenome-assembled genomes of *Idas modiolaeformis* symbionts and used genome-centered metatranscriptomics and metaproteomics to assess key symbiont functions. The Methylophagaceae symbiont is a methylotrophic autotroph, as it encoded and expressed the ribulose monophosphate and Calvin-Benson-Bassham cycle enzymes, particularly RuBisCO. The Nitrincolaceae ASP10-02a symbiont likely fuels its metabolism with nitrogen-rich macromolecules and may provide the holobiont with vitamin B12. The Flavobacteriaceae *Urechidicola* symbionts likely degrade glycans and may remove NO. Our findings indicate that these flexible associations allow for expanding the range of substrates and environmental niches, via new metabolic functions and handoffs.

## Introduction

Obligate chemosynthetic symbioses allow animals and some protists to colonize extreme environments including hydrothermal vents and cold seeps in the deep sea, as well as key productive habitats in coastal areas, such as the seagrass meadows [1, 2]. Chemosynthesis, that is, the assimilation of single-carbon molecules, such as carbon dioxide and methane, using the energy stored in reduced compounds, primarily reduced sulfur (sulfide, thiosulfate, elemental sulfur) and methane, fuels these symbioses. Chemosynthetic symbioses are usually characterized by low diversity and high fidelity of key nutritional symbionts [3]. They exhibit a broad range of transmission strategies, including vertical, horizontal and mixed modes [4], which may determine the fidelity of these associations, as well as the strength of environmental selection for the fittest symbionts and genetic diversity within the symbiont populations [3, 5–7]. As in free-living bacterial populations, genetic diversity broadens the functions of symbionts, allowing the holobiont to acquire new metabolic capabilities and environmental niches [8–10].

Symbiont taxonomic and functional diversity varies markedly in bathymodioline mussels, which include the chemosynthetic genera *Bathymodiolus, Gigantidas* and *Idas* among others [11]. The key symbionts of bathymodioline mussels include the sulfur-oxidizing Thioglobaceae (gammaproteobacterial order PS1, also known as the SUP05 clade), and the *Methyloprofundus* (gammaproteobacterial order Methylococcales) methane-oxidizing bacteria. Some bathymodioline mussels, such as *Bathymodiolus earlougheri, B. billschneideri* and *B. nancyschneideri* host single-species populations of thiothrophic symbionts [12], whereas *Gigantidas childressi* and *G. platifrons* host mainly the methane-oxidizing bacteria [13–15]. Others host both [10, 16]. There are also extreme cases of diversity loss and gain in bathymodioline symbioses. For example, a small deep-sea bathymodioline mussel *Idas argenteus* appears to lack symbionts, which may have been gained in the past and recently lost [17]. Others, such as *Bathymodiolus heckerae*, from the deep Gulf of Mexico, host several taxa of symbionts with distinct functions: apart from two species of methane-oxidizing symbionts and two species of sulfur-oxidizing symbionts, *B. heckerae* has additional bacterial symbionts, including the Methylophagceae sp. methylotrophs, as well as *Cycloclasticus* that catabolize short-chain alkanes [18–20].

Here we focus on another extreme case of multi-species symbioses in bathymodioline mussels namely *Idas* mussels from the Mediterranean Sea. Several previous studies noted the co-occurrence of at least six different gill symbiont phylotypes in an *Idas* species from the east Atlantic and the Mediterranean Sea, This *Idas* species was first identified as *Idas* sp. MED, and is now called *Idas modiolaeformis* [21–24]. These small mussels thrive in seeps and wood falls and the diversity of their symbionts appears to be linked to the availability and composition of reduced fluids [21]. Early studies identified the following six key phylotypes: the *Methyloprofundus sedimenti-related* type I methanotrophs (M1), two phylotypes of Thioglobaceae sulfur-oxidizing symbionts (S1 and S2), Methylophagaceae methylotrophs (M2), as well as two phylotypes that lacked the potential for being chemosynthetic – Bacteroidetes (CFB) and a gammaproteobacterial lineage (G) [23]. Hereafter we refer to the methanotrophs (M1) and thiotrophs (S1 and S2) as the primary, and the others (M2, CFB, and G) as secondary symbionts. Other variants of the secondary symbionts, in particular the G-type, were discovered later [21]. All of these symbionts were found to be associated with bacteriocytes in the symbiont-bearing gill tissue using fluorescence *in situ* hybridization (FISH), however, these observations did not provide conclusive evidence for the intracellular localization of all the symbionts [23]. While the functions of the primary symbionts and the methylotroph M2 were determined based on the detection of marker genes, little is known about the role of the non-chemosynthetic symbiont lineages.

We aimed to characterize the metabolism and physiology of the *Idas* symbionts, using genome-centric metagenomics, metatranscriptomics and metaproteomics. We collected 14 *I. modiolaeformis* individuals from hydrocarbon seeps and brine pools, off the shore of Israel [25, 26]. We hypothesized that the secondary symbionts may contribute to holobiont fitness by providing important functions, based on metabolic handoffs. Alternatively, these potential heterotrophs which are most abundant in mussels that colonize plant debris may extract nutrients from organic substrates [21]. We thus investigated the metabolic potential of the *Idas* symbionts, providing a snapshot of host-symbiont-symbiont interactions in this specific multi-member symbiosis.

## Materials and methods

### Sample collection and processing

Hydrocarbon seeps are found at the toe of Palmahim Disturbance, a large deformation feature at the Israeli margin (Coleman et al., 2011; Gadol et al., 2020) (**Supplementary Fig. S1**). We collected one *Idas* specimen in 2011, from a carbonate sample that was retrieved from the Palmahim pockmark 32°09.6’N and 34°10.0’E at a depth of 1,036 m, using Hercules remotely operated vehicle (ROV), based on the Nautilus E/V. Multiple individuals were collected in 2021 from the edges of a brine pool at the toe of Palamhim disturbance, at 1,150 m water depth (32° 13.4’ N 34° 10.7’ E) [25], using the work-class SAAB Seaeye Leopard ROV, based on R/V Bat Galim (**Supplementary Fig. S1**).

We preserved the individual collected in 2011, as well as 4 individuals that were collected in 2021 at −20°C. DNA was extracted from these individuals using the DNeasy Blood & Tissue Kit (Qiagen), following the manufacturer’s instructions. DNA libraries were constructed for these individuals at HyLabs, Israel, and sequenced as circa 120 million 2×150 bp paired-end reads per sample using Illumina NovaSeq at GENEWIZ, Germany. An additional 4 individuals were preserved in RNAlater. DNA, RNA and proteins were extracted from these individuals using the AllPrep DNA/RNA/Protein Mini Kit (Cat. No. 80004, Qiagen). RNA libraries were constructed for these individuals at Novogene, Singapore, and sequenced as circa 100 million 2×150 bp paired-end reads per sample using Illumina NovaSeq, following library construction with the NEBNext Ultra RNA Library Prep Kit for Illumina (Cat No. 7530) and ribosomal RNA depletion with the Ribo-Zero Plus rRNA Depletion Kit (Bacteria) (Cat No. 20037135) & Ribo-Zero Magnetic Kit (Plant Leaf). For microscopy, 4 additional individuals were fixed in 2% paraformaldehyde in 1x phosphate-buffered saline (PBS) for at most 12 h at 4°C, rinsed three times in 1x PBS, and stored at 4°C in 0.5x PBS/50% ethanol.

### Protein extraction and peptide preparation for metaproteomics

We resuspended the protein precipitates from the 4 individuals extracted with the AllPrep kit in 60 μl of SDT lysis buffer [4% (w/v) SDS, 100 mM Tris-HCl pH 7.6, 0.1 M DTT] and heated to 95°C for 10 min. The SDT protein mixture was cleaned up, reduced, alkylated and digested using the filter-aided sample preparation (FASP) protocol as described previously [27]. We performed all centrifugation steps mentioned below at 14,000 x g. We combined lysates (60 μl) with 400 μl of urea solution (8 M urea in 0.1 M Tris-HCl pH 8.5) and loaded it onto 10 kDa MWCO 500 μl centrifugal filters (VWR International) followed by centrifugation for 20 min. We washed filters once by applying 200 μl of urea solution followed by 20 min of centrifugation to remove any remaining SDS. We performed protein alkylation by adding 100 μl IAA solution (0.05 M iodoacetamide in urea solution) to each filter and incubating for 20 min at room temperature followed by centrifugation for 20 min. The filters were washed three times with 100 μL of urea solution with 15 min centrifugations, followed by a buffer exchange to ABC (50 mM ammonium bicarbonate). Buffer exchange was accomplished by three cycles of adding 100 μl of ABC buffer and centrifuging for 15 min. For tryptic digestion, we added 1 μg of MS grade trypsin (ThermoFisher Scientific) in 40 μl of ABC buffer to each filter and incubated for 16 hours in a wet chamber at 37°C. We eluted tryptic peptides by the addition of 50 μl 0.5 M NaCl and centrifuging for 20 min. Peptide concentrations were determined with the Pierce Micro BCA assay (ThermoFisher Scientific) following the manufacturer’s instructions.

### LC-MS/MS

All proteomic samples were analyzed by 1D-LC-MS/MS as previously described [28]. We loaded 1.2 μg peptide of each sample onto a 5 mm, 300 μm ID C18 Acclaim® PepMap100 pre-column (Thermo Fisher Scientific) with an UltiMateTM 3000 RSLCnano Liquid Chromatograph (Thermo Fisher Scientific) in loading solvent A (2% acetonitrile, 0.05% trifluoroacetic acid). Elution and separation of peptides on the analytical column (75 cm x 75 μm EASY-Spray column packed with PepMap RSLC C18, 2 μm material, Thermo Fisher Scientific; heated to 60°C) was achieved at a flow rate of 300 ml min^-1^ using a 140 min gradient going from 95% buffer A (0.1% formic acid) and 5% buffer B (0.1% formic acid, 80% acetonitrile) to 31% buffer B in 102 min, then to 50% B in 18 min, and finally to 99% B in 1 min and ending with 99% B. The analytical column was connected to a Q Exactive HF hybrid quadrupole-Orbitrap mass spectrometer (Thermo Fisher Scientific) via an Easy-Spray source. Eluting peptides were ionized via electrospray ionization (ESI). Carryover was reduced by a wash run (injection of 20 μl acetonitrile, 99% eluent B) between samples. MS1 spectra were acquired by performing a full MS scan at a resolution of 60,000 on a 380 to 1600 m/z window. MS2 spectra were acquired using data-dependent acquisition selecting for fragmentation the 15 most abundant peptide ions (Top15) from the precursor MS1 spectra. A normalized collision energy of 25 was applied in the HCD cell to generate the peptide fragments for MS2 spectra. Other settings of the data-dependent acquisition included: a maximum injection time of 100 ms, a dynamic exclusion of 25 sec, and the exclusion of ions of +1 charge state from fragmentation. About 50,000 MS/MS spectra were acquired per sample

### Protein identification and quantification

We constructed a protein sequence database for protein identification using the protein sequences predicted from the metagenome-assembled genomes obtained in this study. To identify peptides from the host, we used the annotated protein sequences of the bathymodioline mussel *Bathymodiolus childressi* obtained in a previous study (PRIDE PXD008089-1) [29], since no annotated *Idas* protein sequences were available. We added sequences of common laboratory contaminants by appending the cRAP protein sequence database (http://www.thegpm.org/crap/). The final database contained 49,401 protein sequences and is included in the PRIDE submission (see data access statement) in fasta format. Searches of the MS/MS spectra against this database were performed with the Sequest HT node in Proteome Discoverer 2.3.0.523 as previously described [28]. The peptide false discovery rate (FDR) was calculated using the Percolator node in Proteome Discoverer and only peptides identified at a 5% FDR were retained for protein identification. Proteins were inferred from peptide identifications using the Protein-FDR Validator node in Proteome Discoverer with a target FDR of 5%. To estimate species abundances based on proteinaceous biomass using the metaproteomic data we followed the previously-described approach [30] with the added filter criterion of requiring two protein-unique peptides for a protein to be retained for biomass calculations.

### Bioinformatics

Metagenomes were assembled using SPAdes V3.15 with –meta k=21,33,66,99,127 parameters [31], following adapter trimming and error correction with tadpole.sh, using the BBtools suite following read preparation with the BBtools suite (Bushnell, B, sourceforge.net/projects/bbmap/). Downstream mapping and binning of metagenome-assembled genomes (MAGs) were performed using DAStool, Vamb, Maxbin 2.0 and Metabat2 [32–35] within the Atlas V2.9.1 framework [36], using the genome dereplication nucleotide identity threshold of 0.975. MAG quality was verified using Checkm2 [37] and QUAST [38]. Functional annotation was carried out using the Rapid Annotations using the Subsystems Technology RAST server [39], and key annotations were verified by BLASTing against the NCBI database. RNA reads were quality-trimmed and mapped to the MAGs using BBmap (Bushnell, B, sourceforge.net/projects/bbmap/), and read counts were assigned to coding sequences using FeatureCounts [40]. The counts were normalized as transcripts per million (TPM) [41].

Phylogenetic analyses were performed with IQ-TREE 2 [42] or MEGA11 [43], based on the best model determined by ModelFinder, following sequence alignment with MAFFT [44]. For phylogenomics, we used GTOtree [45] and Fasttree [46], as implemented in GTOtree. Mitochondrial genomes were assembled with MitoZ [47], rotated with MARS [48] and aligned with MAFFT [44].

### Fluorescent in-situ hybridization

Whole tissue from *Idas* individuals was dehydrated, embedded in Steedman’s wax, and cut into 8 μm sections as previously described [20]. To identify the secondary symbionts, we used the Cy3-labeled BhecM2-822, ImedaG-193 and CF319 probes, targeting the Methylophagaceae, Nitrincolaceae and Flavobacteriaceae symbionts, respectively. Cy5-labeled EUB338 targeting most bacteria was used as a positive control (Amann et al., 1990) and the Cy3-labeled NON338 probe was used as a control for background autofluorescence [49]. Photomicrographs were acquired with Nikon A1R Confocal Laser Scanning Microscope, using the Plan Apochromat VC 61.5x objective with oil immersion. The brightness and contrast of the images were adjusted with Adobe Photoshop (Adobe Systems, Inc., USA).

### Data availability

DNA and RNA raw reads, as well as symbiont metagenome-assembled genomes, were deposited to NCBI with project accession number PRJNA930646. Metaproteomic data was submitted to the Proteomics Identification Database PRIDE (accession to be updated, available through the authors).

## Results and discussion

### *Idas modiolaeformis* from the eastern Mediterranean hosts at least six symbiont genotypes

The mussels were identified as *Idas modiolaeformis*, based on the analysis of mitochondrial cytochrome c oxidase I (MT-CO1) sequences from the six metagenomes, using the Barcode of Life Data System identifier. These sequences were 98.4-98.6% similar to those of the *I. modiolaeformis* in the database (for example, the eastern Atlantic individuals, KT216487 in NCBI). The best hits, 98.7-98.9% similarity, were to the sequences of *Idas* individuals recovered from the Nile Deep Sea Fan mud volcano at the depth of 1693 meters (FM212787). Hereafter we refer to our samples as *I. modiolaeformis*, as among other known *Idas* species, the next closest hit was *Idas macdonaldi* with a lower MT-CO1 similarity of 95%. Yet, given that the individuals from the Plamahim Disturbance and the Nile Deep Sea Fan are not genetically identical, it is plausible that genetic exchange between these populations is low. We observed some variation of MT-CO1 and full mitochondrial sequences among the Palmahim individuals corresponding to the occurrence of different haplotypes. Genetic distance may increase with the geographical distance, as the individual that we collected in 2011 from a distinct seep located several hundreds of meters from the brine pool site where other individuals were obtained, was the most diverged (**Supplementary Figure S2**). Yet, as we show later, the host phylotypes were not linked to the symbiont diversity, in agreement with previous findings [21].

We generated metagenome-assembled genomes (MAGs) for the six key *I. modiolaeformis* symbiont phylotypes. We note that besides these six lineages, metagenomic binning resulted in the discovery of MAGs for two additional taxa –Rhizobiales alphaproteobacterium, as well as Bdellovibrionaceae species. Given that the nature of the association between these species and *I. modiolaeformis* is unclear, as they have not been described by amplicon sequence previously, we hereafter focus only on the six previously-detected lineages. For these, good or high-quality genomes were assembled and binned (**Table 1**).

**Table 1:** Taxonomy and completeness statistics for the *Idas modiolaeformis* symbiont metagenome-assembled genomes (MAGs). Compl., completeness; Cont., contamination.

|  | BioSample | Class | Order | Family | Genus | Compl. (%) | Cont. (%) | N50 (Mbp) | Size (Mbp) |
| --- | --- | --- | --- | --- | --- | --- | --- | --- | --- |
| MAG1 | SAMN33052357 | Gammaproteobacteria | Methylococcales | Methylomonadaceae | *Methyloprofundus* | 99.99 | 0.00 | 0.030 | 2.50 |
| MAG2 | SAMN33052358 | Gammaproteobacteria | PS1 | Thioglobaceae | *Thioglobus* A | 90.88 | 0.18 | 0.010 | 1.49 |
| MAG4 | SAMN33052359 | Gammaproteobacteria | PS1 | Thioglobaceae | *Thiodubilierella* | 98.12 | 0.50 | 0.014 | 1.30 |
| MAG4 | SAMN33052360 | Gammaproteobacteria | Nitrosococcales | Methylophagaceae | GCA-002733105 | 99.96 | 0.34 | 0.026 | 2.17 |
| MAG5 | SAMN33052361 | Gammaproteobacteria | Pseudomonadales | Nitrincolaceae | ASP10-02a | 98.18 | 0.01 | 0.037 | 2.10 |
| MAG6 | SAMN33052362 | Bacteroidia | Flavobacteriales | Flavobacteriaceae | *Urechidicola* | 95.79 | 0.30 | 0.020 | 3.36 |
| MAG7 | SAMN33052363 | Alphaproteobacteria | Rhizobiales |  |  | 99.76 | 0.78 | 0.034 | 1.33 |
| MAG8 | NA* | Bdellovibrionia | Bdellovibrionales | Bdellovibrionaceae |  | 73.48 | 1.76 | 0.003 | 1.42 |

\* Not submitted to NCBI due to low completeness

In the previous amplicon-sequencing-based studies the symbionts, particularly the secondary ones, were only classified to class and phylum taxonomic levels. We used our MAGs to obtain improved taxonomic assignments through the Genome Taxonomy Database (GTDB) inference tool and validated their taxonomic placement using phylogenies based on the reference collections of close relative genomes from the GeneBank. We confirmed that the methane-oxidizing symbionts belonged to the *Methyloprofundus* clade, also known as marine methylotrophic group 1 or MMG1, and verified their placement using PmoA protein phylogeny as genomes of most *Methyloprofundus* symbionts are missing from the databases (**Supplementary Fig. S3**). The sulfur oxidizers belong to two distinct clades, *Candidatus* Thioglobus A and *Candidatus* Thiodubilierella (GTDB inference, hereafter *Thioglobus* A and *Thiodubilierella*). The *Thiodubilierella* genus name has been suggested for the symbionts of *B. septemdierum* [50], and was integrated into the GTDB taxonomy. Yet, *Thiomultimodus* has been proposed as an alternative name for both the *Thioglobus* A and *Thiodubilierella* clades, with two SUP05 A and B subclades [10]. Whereas *Thioglobus* A (SUP05 clade A) is often represented by free-living species and clam symbionts, *Thiodubilierella* (SUP05 clade B) has been described only in mussel symbioses [10]. Hereafter we refer only to *Thioglobus* A and *Thiodubilierella* nomenclature for the two thiotrophic symbiont clades. The co-occurrence of these clades in an individual has only been documented in *B. heckerae* [18–20] and *I. modialiformis* [23].

The new MAG-based taxonomic assignments greatly improved the classification of the secondary symbionts and allowed us to make predictions about their physiology based on previously characterized relatives. The methylotrophic symbionts were assigned to the Methylophagaceae clade GCA-002733105. Phylogenic placement based on a tree using single-copy genes indicated that this clade lacks a cultivated representative, and includes lineages commonly found at cold seeps (e.g., the genome of the most closely-related lineage, *Methylophaga* GLR1851 was curated from a sample at the Hikurangi Margin gas hydrate deposits), as well as the only other Methylophagaceae symbionts that occur in *B. heckerae* (**Supplementary Fig. S4**).

Phylotype G belonged to the Nitrincolaceae family ASP10-02a (**Table 1, Supplementary Fig. S5**). Nitrincolaceae are rarely found in symbioses, however, the Rs1 and Rs2 symbionts of the bone-eating worm *Osedax* belong to the Nitrincolaceae family (unnamed genus Rs1). Nitrincolaceae are prominent degraders of nitrogen-rich compounds such as amino acids [51], and the ASP10-02a clade is prominent in the water column, exhibiting often high abundance and activity during algal blooms [52, 53].

The Bacteroidota (CFB) phylotype belonged to the genus *Urechidicola* (Flavobacteriaceae) (**Table 1, Supplementary Fig S6**). The first cultivated *Urechidicola* genus representative *U. process* was isolated from the intestine of a marine spoonworm, *Urechis unicinctus* [54]. Similar to other Flavobacteriaceae, such as the closely-related *Lutibacter, Urechidicola* can degrade multiple organic compounds, including DNA and starch [54, 55]. The closest relative of the *Urechidicola* symbiont in *Idas* was found in a bone-degrading biofilm, and it is represented by GenBank assembly accession GCA_016744415.1. Taxonomic affiliation suggests that both of these secondary symbionts are likely copiotrophs that can catabolize both difficult-to-degrade and protein-rich organic matter such as polysaccharides, nucleic acids and peptides in the marine environment.

The metagenomic read abundance of the primary symbionts varied among the six individuals that were analyzed with metagenomics (**Figure 1**). A single individual (*Idas* 16) lacked the methane-oxidizing and methylotrophic symbionts. In this individual, the Nitrincilaceae symbiont was the most abundant, and *Urechidicola* was found (metatranscriptomics and metaproteomics were, unfortunately, not performed for this individual). We performed metatranscriptomics and metaproteomics on four individuals, which were distinct from the ones used for metagenomics. In these four individuals, the relative abundances of symbionts were consistent between metatranscriptomics and metaproteomics measurements, showing an enrichment of the sulfur oxidizers in sample 3 (5% and 2% biomass, for *Thioglobus* A and *Thiodubilierella*, respectively, **Figure 1, B-D)**. It is important to note here that the relative abundances obtained from metagenomics cannot be compared directly with the relative abundance obtained with metaproteomics, while metagenomics-derived abundances are a good estimator of cell numbers, metaproteomics-derived abundances are a good estimator of species biomass in a microbial population [30]. The methane-oxidizing bacteria were the most abundant in the four samples analyzed with metatranscriptomics (~70-80%) and metaproteomics (~89-98% estimated biomass). FISH analyses substantiated this observation as they showed that the methane-oxidizers were more abundant and larger than the other symbionts (**Figure 2**). The biomass of the other symbionts was much lower in the range of 1.6 to 7.2% for both thiotrophs, with only very few peptides detected for *Urechidicola*. In agreement with the previous study [21], our data suggest that the symbiont proportions differ between individuals, likely based on local environmental conditions experienced by individuals. Methylophagaceae symbionts may require the methane-oxidizers to be present, to supply their substrate methanol. Nitrincolaceae are consistently found among the samples, with both the meta-omics and FISH analyses (**Figure 2**) and are metabolically active, as their genes and proteins were expressed. Methylophagaceaeae, Nitrincolaceae and Urechidicola symbionts appear to co-occur with the primary symbionts in the gill tissue where the primary symbionts are present (**Figure 2**), in agreement with previous data [23]. Yet, it is still not clear if the secondary symbionts are extra- or intra-cellular.

**Figure 1:**
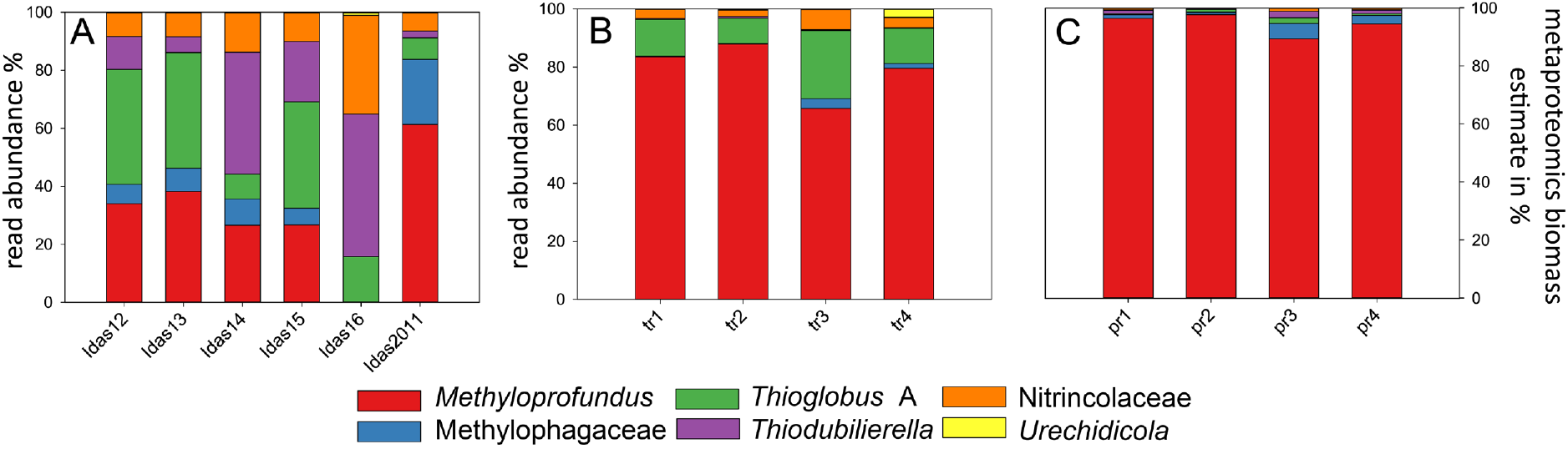
The relative abundance and biomass distribution of the six key symbiont genotypes in individual *Idas modiolaeformis* specimens: (A) read abundance in metagenomes (Idas12-16 are samples from the brine pool site, Idas2011 has been collected in 2011 from carbonates at a nearby seep site), (B) read abundance in metatranscriptomes, (C) biomass estimates based on proteomics following Kleiner et al. 2017. Metatranscriptomics and metaproteomics were performed on the same four individuals (tr1-4 are equivalent to pr1-4). Due to the low confidence of detection, *Urichidicola* proteins were not counted in panel C.

**Figure 2:**
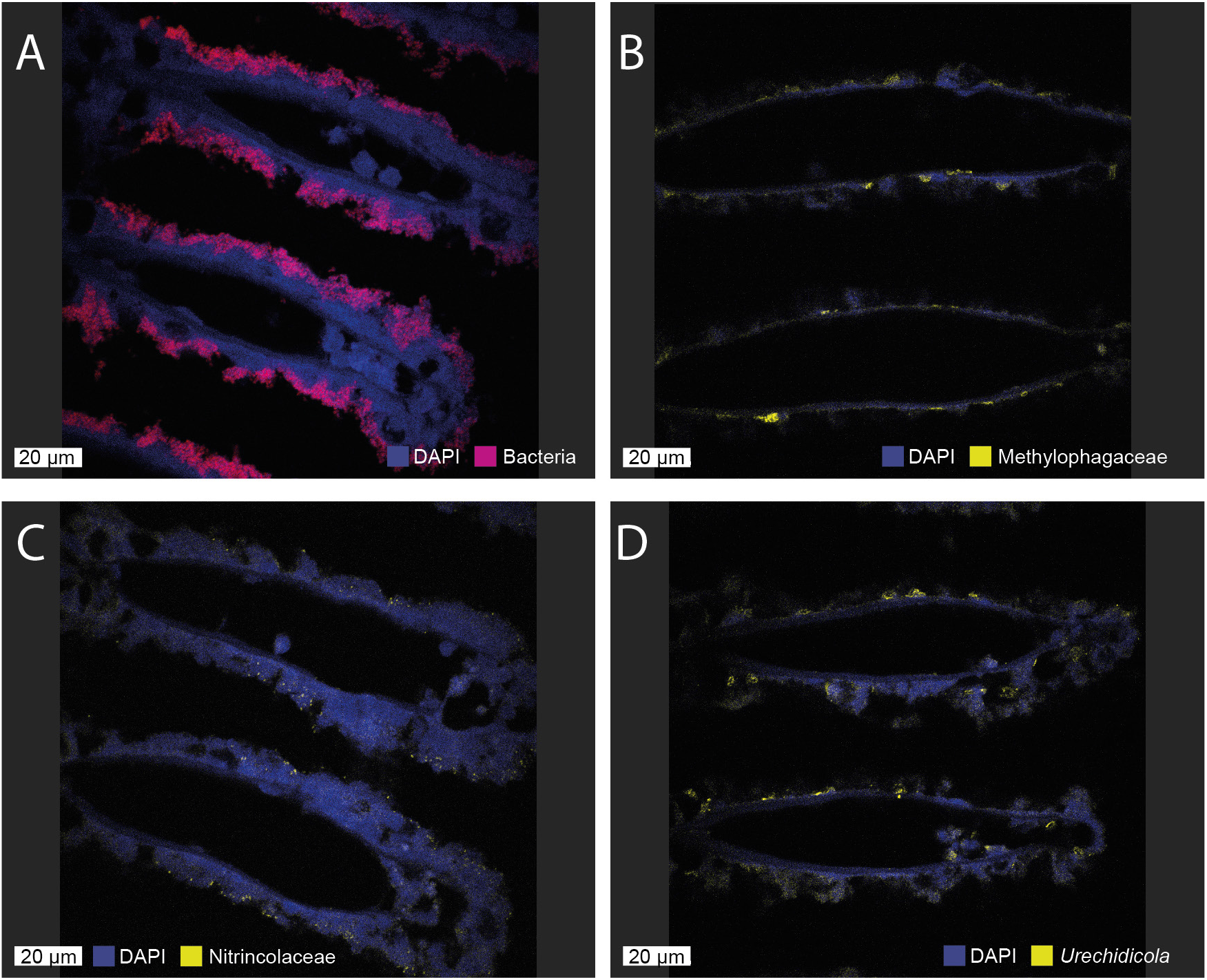
Fluorescent in situ hybridization (FISH) shows the presence of the secondary symbionts in the *Idas* gill tissue. A) All the symbiotic bacteria are detected with the EUB338 probe (red), and large methanotroph morphotypes can be observed. B) Methylophagaceae symbionts (BhecM2-822 probe, yellow) appear to often aggregate along the gill tissue. C) Nitrincolaceae symbionts (ImedaG-193 probe, yellow) were found sporadically throughout the tissue. D) Flavobacteriales (*Urechidicola*) symbionts in the gill tissues (CF319, yellow). DAPI staining was used to visualize all DNA. The scale bar is 20 μm.

### Oxidation of methane and its products fuel the *Idas* symbiosis

Methane oxidizers are the key nutritional symbionts of *I. modiolaeformis*, as suggested by their abundance in the metatranscriptomes and metaproteomes. Additionally, we performed carbon stable isotope analysis of *Idas* individuals that showed a very light carbon isotope ratio of - 58‰ δ^13^C vs. Vienna Pee Dee Belemnite (Zvi-Kedem et al., in prep.), which indicates that the majority of carbon in *Idas* is derived from the isotopically light methane. The particulate methane monooxygenase PmoCAB complex is most well-expressed at both mRNA and protein levels (**Supplementary Fig. S7, here and hereafter see Supplementary Table S1 for the full list of genes and proteins expressed**). The lanthanide-containing XoxF type methanol dehydrogenase is the predominant one, whereas the calcium-dependent enzyme MxaFI was found, but expressed at lower levels. Carbon assimilation and energy conservation from formaldehyde oxidation function via the ribulose monophosphate (RuMP) pathway, using only the Entner–Doudoroff (ED) bypass, but not the Embden–Meyerhof–Parnas (EMP) variant [56], as the *pfk* gene encoding the pyrophosphate-dependent phosphofructokinase has not been found in the genomes of the *Methyloprofundus* symbiont of *I. modiolaeformis* (**Supplementary Fig. S7**). This is similar to *Bathymodiolus japonicus* symbionts and in contrast to those of *Bathymodiolus platifrons*, which have both the ED and EMP variants [57]. The serine cycle is incomplete, as hydroxypyruvate reductase is missing, however, some central proteins of this pathway, such as the serine--glyoxylate aminotransferase and malyl-CoA lyase, were substantially expressed at both the RNA and amino acid levels, indicating that the partial serine pathway is functional (**Supplementary Fig. S7**) [16]. All the genes in the tricarboxylic acid (TCA, Krebs) cycle were found and expressed, therefore energy can be conserved through the oxidation of compounds derived from methane assimilation, such as pyruvate (**Supplementary Fig. S7**). Formaldehyde detoxification and oxidation to CO_2_ via the dissimilatory tetrahydromethanopterin route is likely prominent, due to the high expression of this pathway, in particular, that of the 5,6,7,8-tetrahydromethanopterin hydro-lyase (*fae* gene, **Supplementary Fig. S7**). Both oxygen and nitrate respiration is feasible, given the presence of the respiratory NarGHIJ and NirK, yet the respiratory complex IV is expressed at much higher levels, indicating that oxygen is the key electron acceptor for energy conservation (**Supplementary Fig. S7**).

The metabolism of the Methylophagaceae symbiont is a rare case of methylotrophic autotrophy in chemosynthetic symbioses. Their co-occurrence with the methane oxidizers suggests that they might use methanol, formaldehyde and other metabolites produced by the methane oxidizers. Methane oxidizers excrete metabolites such as methanol, formaldehyde, formate, acetate, and succinate [58–60], which can be used by co-occurring organisms, in particular, methylotrophs [61]. Partnerships between methane oxidizers and methylotrophs are widespread in free-living communities [61, 62], and our results suggest that this association plays a role in *Idas* symbiosis.

The Methylophagaceae symbionts lack methane monooxygenases, yet highly express the pyrroloquinoline quinone (PQQ)-dependent methanol dehydrogenase, as well as the key enzymes of the RuMP pathway for formaldehyde assimilation (**Figure 3**). As opposed to the primary methane-oxidizing symbionts, the methylotrophs encoded and expressed both the ATP-dependent EMP and ED variants of the RuMP pathway, which have distinct energetic demands [56]. Formate oxidation is also more flexible in the methylotrophs, as we identified the occurrence and expression of not only the two-subunit tungsten-containing formate dehydrogenase FdhAB, but also the respiratory molybdoenzyme dehydrogenase-O (FdoGHI) that catalyzes the oxidation of formate to carbon dioxide, donating electrons to the membrane soluble quinone pool [63–66]. This hints that the oxidation of formate alone can fuel the metabolism of the Methylophagaceae symbiont. In addition to originating from methane oxidation, the used formate could also come from mitochondrial metabolism [67]. In terms of terminal electron acceptors (TEA) for methanol and formate oxidation we only found genes for the use of oxygen as TEA and no genes for other TEAs such as nitrate.

**Figure 3:**
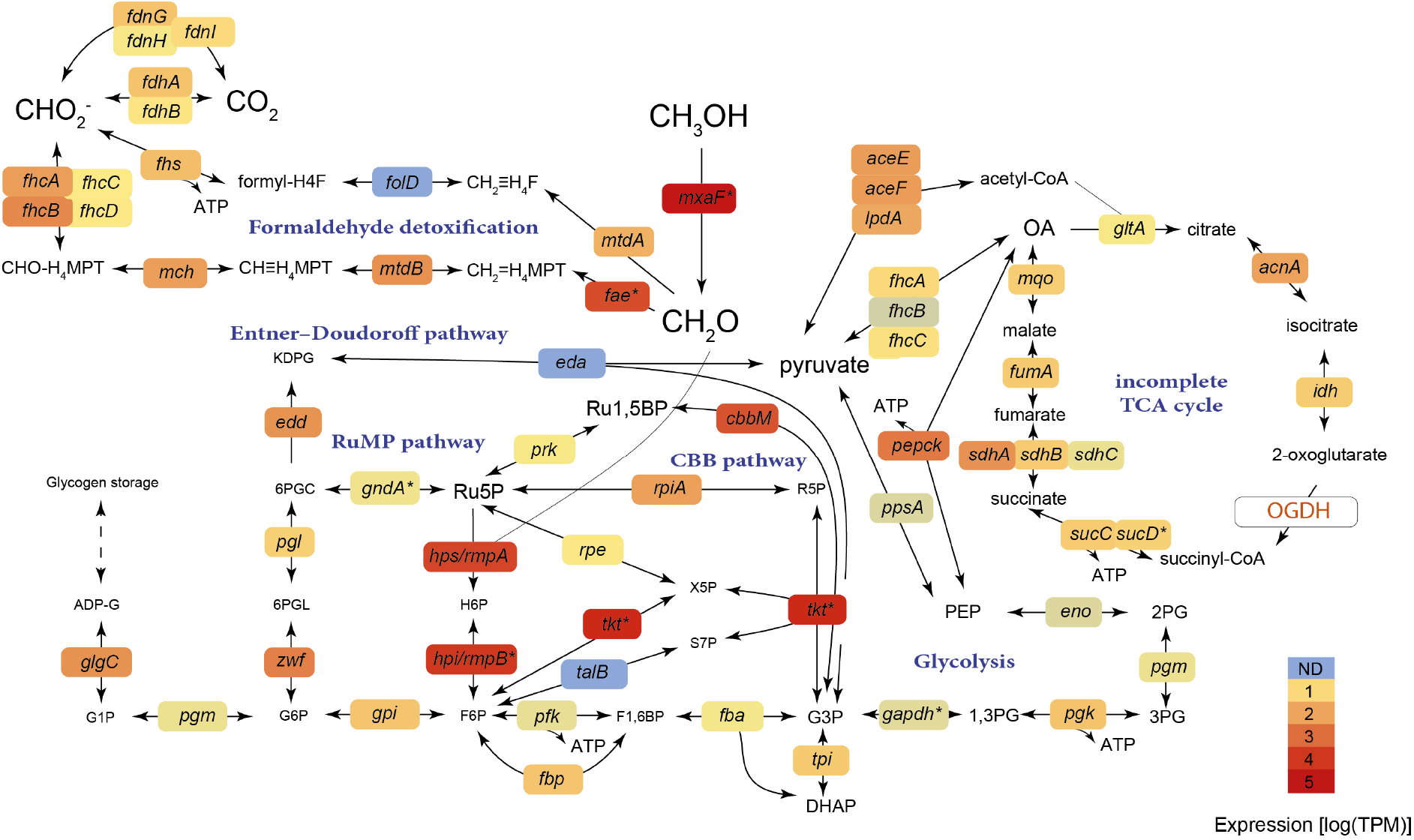
Central carbon metabolism in the Methylophagaceae symbiont of *Idas modiolaeformis*. Both the Embden–Meyerhof–Parnas (EMP) and Entner–Doudoroff (ED) variants of the ribulose monophosphate (RuMP) pathway are feasible. This bacterium can fix inorganic carbon via the Calvin–Benson-Bassham (CBB) cycle. The tricarboxylic acid cycle is incomplete, as the 2-oxoglutarate dehydrogenase (OGDH) was not found. The genes are as follows: methanol dehydrogenase *mxaF*; 3-hexulose-6-phosphate synthase *hps/rmpA*; 3-hexulose-6-phosphate isomerase *hpi/rmpB*; transketolase *tkt*, ribose-5-phosphate isomerase *rpiA*; phosphoribulokinase *prk*; ribulose-phosphate 3-epimerase *rpe*; transaldolase *talB*; ATP-dependent 6-phosphofructokinase *pfk*; fructose-1,6-bisphosphate aldolase/phosphatase *fbp*: glucose-6-phosphate isomerase *gpi*; glucose-6-phosphate 1-dehydrogenase *zwf*; 6-phosphogluconolactonase *pgl*; phosphogluconate dehydratase *edd*; 2-dehydro-3-deoxy-phosphogluconate/2-dehydro-3-deoxy-6-phosphogalactonate aldolase *eda*; phosphoglucomutase *pgm*; glucose-1-phosphate adenylyltransferase *glgC*; fuctose-bisphosphate aldolase *fba*; triosephosphate isomerase *tpi*; formaldehyde activating enzyme *fae*; methylene tetrahydromethanopterin dehydrogenase *mtdB*; methenyltetrahydromethanopterin cyclohydrolase *mch*; formyltransferase/hydrolase complex *fhcABCD*; NAD(P)-dependent methylenetetrahydromethanopterin dehydrogenase *mtdA*; bifunctional methylenetetrahydrofolate dehydrogenase / methenyltetrahydrofolate cyclohydrolase *fold*; formate--tetrahydrofolate ligase *fhs*; formate dehydrogenase *fdhAB*; formate dehydrogenase, nitrate-inducible *fdnGHI*; glyceraldehyde-3-phosphate dehydrogenase *gapdh*; phosphoglycerate kinase *pgk*; phosphoglucomutase *pgm*; enolase *eno*; phosphoenolpyruvate synthase *ppsA*; phosphoenolpyruvate carboxykinase *pepck*; oxaloacetate decarboxylase Na(+) pump *oadABC*; pyruvate dehydrogenase *aceEF-lpdA*; citrate synthase *gltA*; aconitase *acnA*; isocitrate dehydrogenase *idh*; 2-oxoglutarate dehydrogenase complex *OGDH*; succinate--CoA ligase *sucCD*; succinate dehydrogenase *sdhABC*; fumarate hydratase class I, aerobic *fumA*; malate:quinone oxidoreductase *mqo*. Metabolites: OA, oxaloacetate; PEP, phosphoenolpyruvate; 2-phosphoglycerate, 2PG; 3-phosphoglycerate, 3PG; 1,3-bisphosphoglycerate 1,3BPG; 3-phosphoglyceraldehyde, G3P; dihydroxyacetone phosphate, DHAP; fructose 1,6-bisphosphate, F1,6BP; fuctose 6-phosphate, F6P; hexulose 6-phosphate, H6P; ribulose 5-phosphate, Ru5P; ribulose-1,5-bisphosphate, Ru1,5BP; ribose 5-phosphate, R5P; glucose 6-phosphate, G6P; 6-phosphogluconolactonase, 6PGL; 2-Dehydro-3-deoxy-D-gluconate 6-phosphate, 6PGC; 2-keto-3-deoxy-6-phosphogluconate, KPDG; gucose 1-Phosphate, G1P; ADP-glucose, ADP-G; tetrahydrofolate, H4F; tetrahydromethanopterin, H4MPT. Average expression values from 4 individuals are shown.

Alongside the RuMP pathway, Methylophagaceae symbionts have the genes for all steps of the Calvin–Benson–Bassham (CBB) cycle and highly express the key enzyme, form II ribulose-1,5-bisphosphate carboxylase/oxygenase (RuBisCO, **Figure 3**). We note that although form I RuBisCo appear to be widespread in Methylophagaceae, form II RuBisCo is rare in this clade (**Supplementary Fig. S7**). These closely related RuBisCo forms differ in their affinity to oxygen and CO_2_, and form II enzymes have a low specificity factor, that is, function better under high CO_2_ and low O_2_ conditions [68–70]. Such conditions may exist in the host’s bacteriocytes, giving an advantage to bacteria with form II RuBisCO [71].

Similar to obligate autotrophs [72, 73], the TCA cycle appears to be incomplete in most cultivated Methylophagaceae, which lack the 2-oxoglutarate dehydrogenase activity [74–76]. We did not find the 2-oxoglutarate dehydrogenase-encoding genes in the high-quality MAG of the symbiotic Methylophagaceae, indicating that the TCA cycle is incomplete. The incomplete TCA allows for the production of intermediates from one-carbon (C1) compounds and prevents futile cycling, that is, the destruction of larger organic compounds via catabolism. This is typical of obligate methylotrophs, such as *Methylobacillus flagellates* [77].

### Variability in terminal electron acceptors, metabolite uptake and vitamin usage may lead to niche differentiation among the thiotrophic symbionts

The two sulfur-oxidizing symbionts, *Thioglobus* and *Thiodubilierella* species, appear to be obligate autotrophs as the key enzymes of the TCA cycle, 2-oxoglutarate dehydrogenase and malate dehydrogenase, were not found in the MAG (**Supplementary Figure S8**). This is in agreement with previous observations of symbiotic and free-living organisms from this clade [10, 78, 79]. Both sulfur-oxidizing symbionts have the genes to gain energy from sulfur compounds using the sulfide: quinone oxidoreductase (Sqr), the Sox sulfur oxidation system (SoxYZXAB) and the reverse dissimilatory sulfite reduction (rDSR) system, which catalyzes the oxidation of sulfide to sulfate via the adenosine-5’-phosphosulfate, using the adenylylsulfate reductase and sulfate adenylyltransferase for the two final steps of the pathway. The genes encoding these functions were among the most abundantly transcribed ones in the metatranscriptomes and were often detected at the protein level (**Supplementary Figure S8**). Both lineages highly expressed the Calvin-Benson-Bassham cycle enzymes, in particular the two subunits of the type I RuBisCo (in particular, the *rbclLS* genes and respective proteins). These obligate autotrophs likely can import some organic compounds similar to the symbionts of *Bathymodiolus azoricus* [16], as TRAP-type dicarboxylate transport systems were encoded and expressed (C4-dicarboxylate and unknown substrate 6 in *Thioglobus*, only the latter in *Thiodubilierella*).

Apart from the cytoplasmic assimilatory nitrate reductase, only *Thiodubilierella* carries the periplasmic NapABGH, confirming the modularity of nitrogen metabolism in the Thioglobaceae clade [10]. Some discrepancies in oxygen respiration were found: *Thioglobus* encoded and highly expressed only the *ccoNOP* genes encoding the cbb_3_-type cytochrome c oxidase, whereas *Thiodubilierella* encoded three variants (cbb_3_, bo_3_, ba_3_). These terminal oxidases differ in their affinity to O_2_ and H_2_S [80], likely highlighting adaptation to distinct redox conditions. Another key difference could be the dependence on cobalamin (vitamin B12, see below). These variations may determine the selection and thus relative abundance of *Thioglobus* and *Thiodubilierella* symbionts.

### Secondary symbionts are heterotrophs that can use nitrogen-rich and difficult-to-degrade metabolites

All three meta-omics suggest that the Nitrincolaceae symbiont is a heterotroph that is capable of using numerous substrates, given the presence and substantial expression of the complete TCA cycle, as well as that of multiple transport systems for organic compounds, such as peptides, amino acids and nucleosides. Relevant examples include: (1) most enzymes for the degradation of branched-chain amino acids co-occurred in the genomes and were transcribed (**Figure 4**); (2) a cluster 11 RfuABCD riboflavin/purine/nucleoside transporter, which clustered with nucleoside degradation enzymes, such as deaminases and phosphorylases (Cda, Ada, DeoABC); (3) the putrescine transport (PotABCD) and degradation system (PuuABCDE), although the *puuC* gene was not found, likely due to an assembly issue, as these genes occurred at the edge of a contig. An additional gene cluster encoded the complete PaaA-K and HpaA-I machinery for the catabolism of phenylacetate and hydroxyphenylacetate [81]. This pathway allows for the degradation of recalcitrant, plant-derived organic aromatics, especially in marine bacteria that thrive under fluctuating oxygen conditions in the Mediterranean Sea [82, 83]. Some catabolic reactions may be carried out anaerobically, particularly given the fermentative potential of this bacterium, suggested by the presence of lactate dehydrogenases (some expressed at the mRNA level, **Figure 4**). We also identified the genes for nitrate respiration to nitric oxide including the genes for the respiratory nitrate reductase NarGHIJ and copper-dependant nitrite reductase NirK, and *nirK* was among the most well-expressed genes at the mRNA level in this symbiont (**Figure 4**). However, oxygen is likely the key electron acceptor for these symbionts as the cbb_3_- and caa_3_ complex IV-encoding genes were highly expressed at the mRNA level (**Figure 4**).

**Figure 4:**
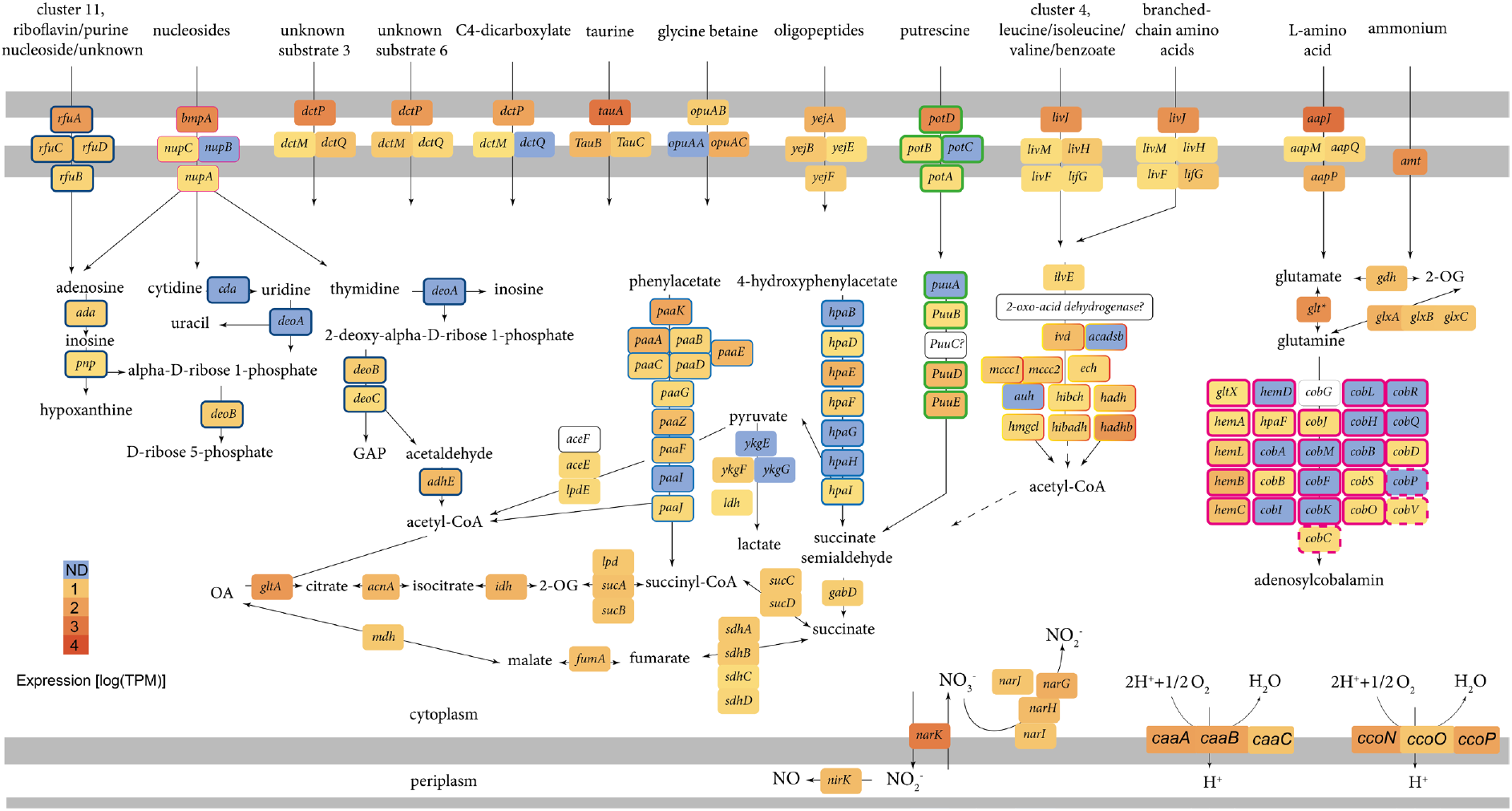
The Nitrincolaceae symbiont can import and catabolize multiple metabolites, including peptides, amino acids (in particular, branched-chain amino acids), nucleosides and some aromatics, such as phenylacetate. The tricarboxylic acid cycle is complete. The symbiont can likely ferment pyruvate to lactate, as the cytochrome c-dependent D-lactate and L-lactate dehydrogenases were found and expressed. A near-complete pathway of adenosylcobalamin (B12) is encoded and partially transcribed. * indicates detection at the protein level. Frames of the same color indicate gene clusters. The following genes are shown: adenosine deaminase *ada*, polyribonucleotide nucleotidyltransferase *pnp*; cytidine deaminase *cda*; thymidine phosphorylase *deoA*; phosphopentomutase *deoB*; deoxyribose-phosphate aldolase *deoC*; aldehyde-alcohol dehydrogenase *adhE*; acetyl-CoA carboxylase *accEF-lpdE*; L-lactate dehydrogenase *ykgEFG*; L-lactate dehydrogenase *ldh*; citrate synthase *gltA*; aconitase *acnA*; isocitrate dehydrogenase *idh*; 2-oxoglutarate dehydrogenase complex *sucAB-lpdE*; succinate--CoA ligase *sucCD*; succinate dehydrogenase *sdhABC*; fumarate hydratase class I, aerobic *fumA*; malate dehydrogenase *mdh*; pyruvate dehydrogenase *aceEF-lpdA*; phenylacetate catabolon *paaA-K*; 4-hidroxyphenylacetate catabolon *hpaB-I*; succinate-semialdehyde dehydrogenase [NADP(^+^)] *gabD*; putrescine catabolon *puuA-D*; acetolactate synthase *ilvB*; isovaleryl-CoA dehydrogenase *ivd*; short/branched chain specific acyl-CoA dehydrogenase *acadsb*; enoyl-CoA hydratase *ech*; methylcrotonoyl-CoA carboxylase *mcc1,2*; methylglutaconyl-CoA hydratase *auh*; 3-hydroxyisobutyryl-CoA hydrolase *hibch*; hydroxyacyl-coenzyme A dehydrogenase *hadH*; hydroxymethylglutaryl-CoA lyase *hmgcl*; 3-hydroxyisobutyrate dehydrogenase *hibadh*; acetyl-CoA acyltransferase *hadhb*; glutamine synthetase *glt*; glutamate dehydrogenase *gdhA*; glutamine amidotransferase *glxABC*; cbb_3_-type cytochrome c oxidase *ccoNOP*; caa_3_-type cytochrome c oxidase *ccaABC*; respiratory nitrate reductase *narGHIJ*, copper-containing nitrite reductase *nirK*; nitrate/nitrite antiporter *narK*;. The putative transporter substrates are mentioned in the figure. Glyceraldehyde 3-phosphate, GAP; oxaloacetate, OA; 2-oxoglutarate, 2-OG. Average expression values from 4 individuals are shown.

A key feature of the metabolism encoded in the *Urechidicola* symbiont genome (Flavobacteriaceae, Bacteroidota) is its potential to degrade glycans (polysaccharides). Bacteroidota, Flavobacteriaceae in particular, are ubiquitous degraders of complex glycans, found in many environments from human gut mucus to algal blooms in the ocean [84–87]. The diversity of glycans is very large and their degradation is carried out by an array of carbohydrate-active enzymes (CAZymes) that are often organized in polysaccharide utilization loci (PULs), which are typical of Bacteroidota [84]. The key features of PULs are the presence of SusCD transporters, which comprise the cell surface glycan-binding lipoprotein SusD and the outer membrane TonB-dependent transporter SusC, which can be found in multiple copies in Bacteroidota genomes [88, 89]. We identified four SusCD pairs in the *Urechidicola* symbiont, two of which were among the top 50 abundantly transcribed genes in the metatranscriptomes, and one SusC protein was detected in the metaproteomes despite the overall protein identification rate for this symbiont. We note that genomic coding sequences’ coverage was very low for *Urechidicola* in the metatranscriptome (17%) and metaproteome (5%, mostly with low confidence), and only the most active features are highlighted by these analyses. Given the considerable fragmentation of this genome (300 scaffolds), we couldn’t identify large PULs, yet multiple CAZymes were found and often clustered (**Figure 5**). For example, a single 30996 bp long contig contained two SusCD pairs, as well as multiple CAZymes, such as DD-carboxypeptidase EC 3.4.16.4; L-Ala-D/L-Glu epimerase EC 5.1.1.20; glucosamine-6-phosphate deaminase EC 3.5.99.6; N-acetylmuramic acid 6-phosphate etherase EC 4.2.1.126; as well as AmpG family muropeptide MFS transporter; glycosyl hydrolases of family 18; family 10, and family 3 (**Figure 5)**. Thus, genomic and transcriptomic data provide evidence for the glycan degrading activity of *Urechidicola*, yet whether the substrates are derived from the gill (e.g mucus), or the environment remains unknown. To date, FISH analyses do not show clearly if these bacteria are endo- or ectosymbionts of *Idas* gills. One piece of evidence suggesting that they might be ectosymbionts is the fact that cell adhesion appears to play an important role for *Urechidicola*, as a sequence encoding the FAS1 (fascilicin) domain protein is among the top 20 most abundantly transcribed genes [90, 91]. One hypothesis that remains to be explored is that there are metabolic handoffs between *Urechidicola* and Nitrincolaceae symbionts, for example, via the exchange of aromatic deriviates of polymer degradation, such as phenylacetate and hydroxyphenylacetate [92].

**Figure 5:**
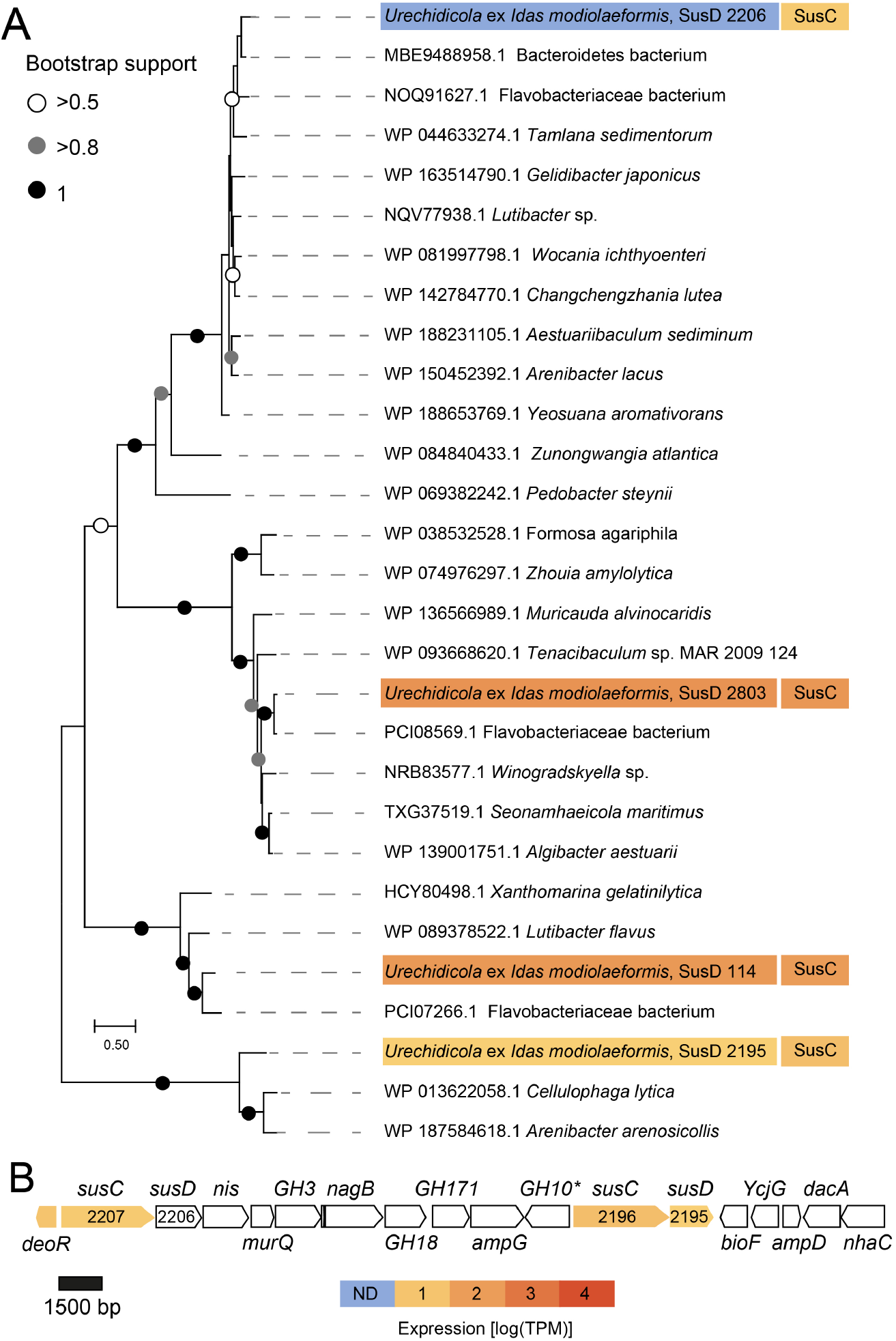
Polysaccharide utilization loci (PULs) in *Urechidicola* symbionts of *Idas modiolaeformis*. A) Phylogeny and mRNA-level expression of the SusD proteins (MEGA11, 576 amino acid positions, LG+G+I model). The expression values are based only on a library from a single *Idas* individual, in which expression of these rare symbionts had detectible coverage. B) The largest contiguous PUL (~27000 bp) in *Urechidicola* symbiont comprised two *susCD* pairs, as well as genes encoding the following: transcription regulator deoR; sodium/iodide co-transporter, *nis*; N-acetylmuramic acid 6-phosphate etherase, *murQ*, glucosamine-6-phosphate deaminase, *nagB*; MFS permease, *ampG*; aminotransferase, *bioF*; L-Ala-D/L-Glu epimerase, *ycgG*; N-acetylmuramoyl-L-alanine amidase, *ampD*; D-Ala-D-Ala carboxypeptidase, *dacA*; Na+/substrate antiporter, *nhaC*; as well as glycoside hydrolases 3,18,171,10. Most of these genes are involved in murein metabolism. * Was identified by NCBI domain search, but not by dbCAN. The full list of CAZymes is available in **Supplementary Table S2**. The tree scale represents the number of substitutions per site. Expression values from a single individual (TR4, detectible *Urechidicola* coverage) are shown.

### Nitrincolaceae symbionts may contribute to holobiont fitness by producing vitamin B12

The Nitrincolaceae symbiont can produce adenosylcobalamin (vitamin B12), given the presence of almost all the genes needed for its synthesis (**Figure 4**). The only gene out of a total of 25 for vitamin B12 biosynthesis that we did not find is the *cobG* gene, which is likely due to the incompleteness of the MAG. Many of the vitamin B12 biosynthesis genes were transcribed and detected in the metatranscriptomes. We also found the genes for vitamin B12 biosynthesis in the methane-oxidizing symbiont, which is a likely key producer of adenosylcobalamin in *I. modiolaeformis*, given its dominant biomass in most individuals (**Figure 1**). In hosts that lack the methane-oxidizing symbionts, B12 production by the alternative Nitrincolaceceae symbionts may be crucial for the holobiont. *Nitrincolaceae*, specifically the ASP10-02a lineage to which the symbionts belong, are i) dominant species in Earth’s cold oceans, ii) are often associated with algal bloom degradation, iii) appear to be the most prominent B12 producers in these habitats and iv) supply B12 specifically to Methylophagaceae, suggesting that B12-based ASP10-02a-methylotroph/autotroph associations can be advantageous under some conditions [53, 93]. The B12-based dynamics play a key role in the water column [94], but also human gut [95] and insect symbioses [96], and thus may be crucial also for chemosynthetic symbioses. The outer membrane receptor BtuB of the vitamin B12 transporter was found in the genomes *Methyloprofundus*, *Thioglobus_A, Urechidicola* and Methylophagaceae symbionts. Whereas methane-oxidizing symbionts are likely prototrophs for cobalamin, but still can take it up from the environment, the other symbionts are auxotrophs and may depend on its uptake.

In bacteria, the key reaction that may depend on cobalamin is methionine synthesis, as methionine synthase, thus the lack of this vitamin may hinder DNA synthesis, methionine regeneration and lead to homocysteine accumulation [97]*. Thiodubilierella* appears to lack the *btuB* gene needed for B12 import, however, unlike the *Thioglobus* A symbiont, which encodes only the cobalamin-dependent methionine synthase (MetH; EC 2.1.1.13), *Thiodubilierella* has both the MetH, and the cobalamin-independent methionine synthase (MetE; 5-methyltetrahydropteroyltriglutamate–homocysteine methyltransferase; EC 2.1.1.14), and therefore may not depend on cobalamin for production of the essential amino acid methionine and thus growth [96]. Similar to insect symbioses [96], the interplay between cobalamin requirement, synthesis and uptake may contribute to determining the complexity of *Idas* symbiosis. In our metagenomic dataset, the realtiove abundances of Nitrincolaceae and *Thiodubilierella*, but not Nitrincolaceae and Thioglobus_A symbionts appear to positively correlate (R^2^=0.79 and 0.03 for linear regression of centered-log transformed data). Although compositional data correlation should be treated with caution, our results hint at the interdependence of these two symbiont populations.

### *Urechidicola* symbionts may denitrify to N_2_, removing NO

*Urechidicola* appears to be the only bacterium capable of complete denitrification to N_2_. Nitrate and nitrite reduction can be catalyzed by the periplasmic enzymes heterodimeric nitrate reductase (NapAB, no expression found) and the NrfAB cytochrome c552 nitrite reductase (both subunits were expressed at the mRNA level). The *norBC* genes that encode the respiratory nitric oxide (NO) reductase were highly transcribed. Most importantly, the *nosZ* gene encoding the periplasmic sec-dependant nitrous oxide reductase was the 11^th^ most abundantly transcribed gene in this bacterium. Both the methane-oxidizing and methylotrophic symbionts can oxidize nitrite to nitric oxide but lack the genes needed to complete the remaining steps of denitrification. Symbiont nitrate respiration can contribute to the holobiont fitness, as it fuels energy conservation during hypoxia induced by natural perturbations or when the shell valves are closed, and reduces competition between host and symbiont for oxygen [16]. Removal of toxic nitrite respiration product NO could benefit the symbionts [98]. Whereas the *Urechidicola* symbiont is only present in low abundance, complete denitrification is likely beneficial mainly for their population, which can take advantage of excess NO produced by the host or the other symbionts, but in some cases may also contribute to NO removal at the holobiont level.

## Conclusions

*The I. modialiformis* symbiosis has a higher number of symbiont species than the typical chemosynthetic associations in most other hosts. The metabolic flexibility in these symbioses expands the range of catabolized substrates and likely allows for the colonization of not only chemosynthetic environments but also organic substrates, such as wood. This nutritional plasticity may have played a role in the adaptation and evolutionary transition of bathymodioline mussels from organic substrates to the deep sea [99]. Expanding the symbiont diversity and substrate range may lead to energetic costs, resulting in lower growth rates or reduced reproductive output [21]. These costs may be reduced as only the symbiont that can use the locally abundant substrate grows up to high abundances. For example, at seeps and brine pools methane fuels the holobiont metabolism, as the methanotrophic symbiont dominates in terms of biomass, and due to the very light carbon stable isotope signature. Even in the absence of relevant amounts of external organic substrates the secondary symbionts might provide additional benefits which include the recycling, and efficient use of resources such as NO, methanol and formate, as well as sharing goods such as vitamin B12 (**Figure 6**). Some of these positive interactions have equivalents in free-living communities, suggesting that the complex symbiotic associations may have evolved based on the existing interactions.

**Figure 6:**
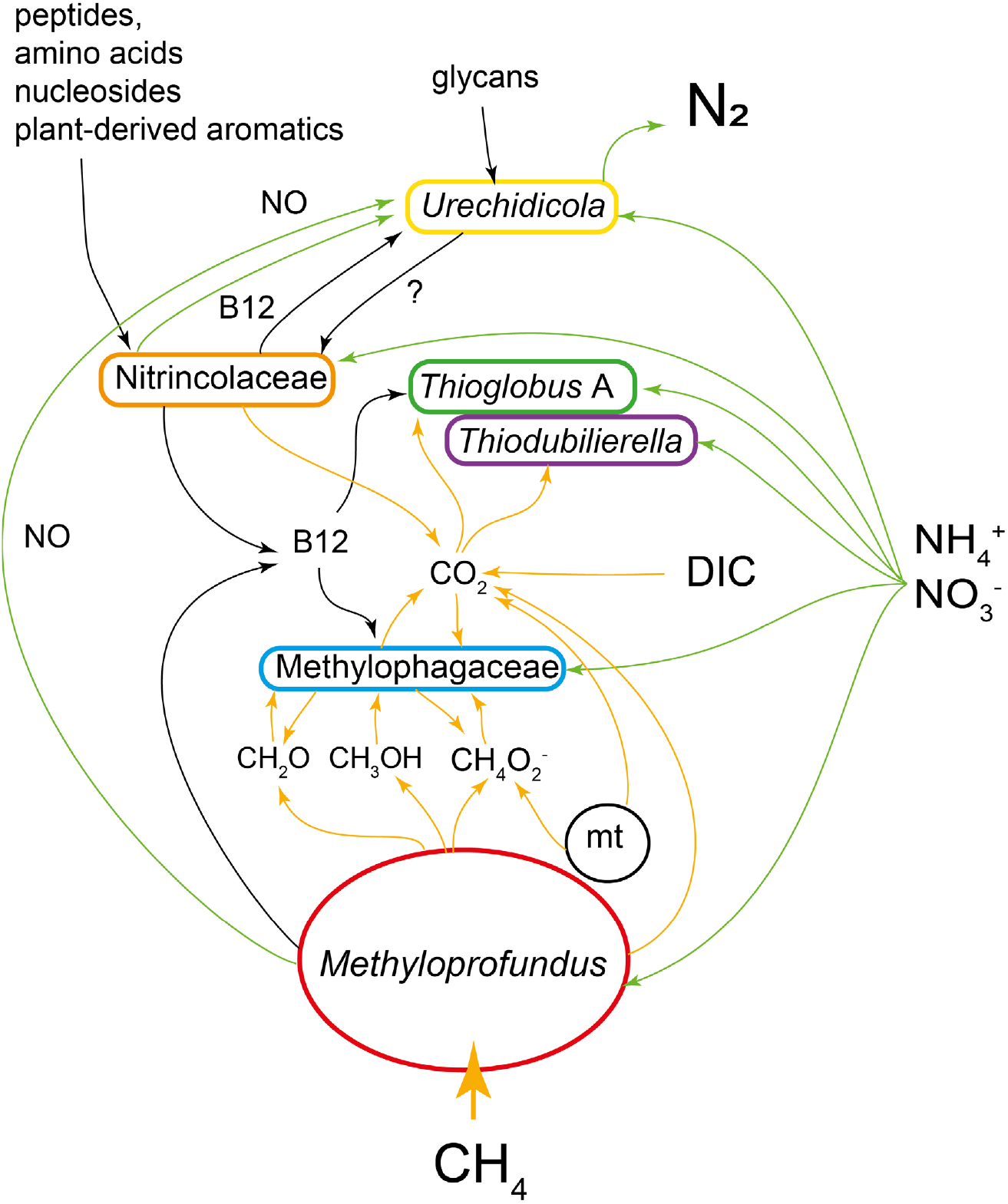
Schematic representation of key metabolic handoffs in the *Idas* symbioses. C1 routes (yellow), nitrogen (green) and macromolecule (black) are highlighted by arrows. For nitrate, the routes comprise both assimilatory and dissimilatory ones. A mitochondrion is included (mt). DIC is dissolved inorganic carbon.

## Author contributions

T.Z-K, M. R-B and D.T. conceived this study. T.Z-K conducted microscopy. M. R-B. performed bioinformatics. S.V. and M.K. performed proteomics. M.R-B and T.Z-K. compiled the data. M.R-B, D.T. and M.K. acquired funding. T. Z-K, M. R-B and M.K. wrote the paper with the contributions of all co-authors.

## Acknowledgments

The authors thank all individuals who helped during the expeditions, including onboard technical and scientific personnel, the captains and crew of the R/V Bat Galim and E/V Nautilus, and teams operating ROVs “Hercules” and “Yona”. This research used samples and data provided by the E/V Nautilus Exploration Program - expedition NA019. We thank Sharon Tal at the Histology Service Unit at the University of Haifa and Boris Shklyar at the Bioimaging unit, Faculty of the Natural Sciences, University of Haifa.

## Funding

This study was funded by the Israeli Science Foundation (ISF, grant 913/19 to M. R-B), the U.S.-Israel Binational Science Foundation (BSF, grant no. 2019055 to M. R.-B. and M. K.), the Israel Ministry of Energy (grant no. 221-17-002 to M. R-B) and the US National Science Foundation (grant IOS 2003107 to M. K.).

## Conflicts of interest

None declared.

## Supplement

**Figure S1:**
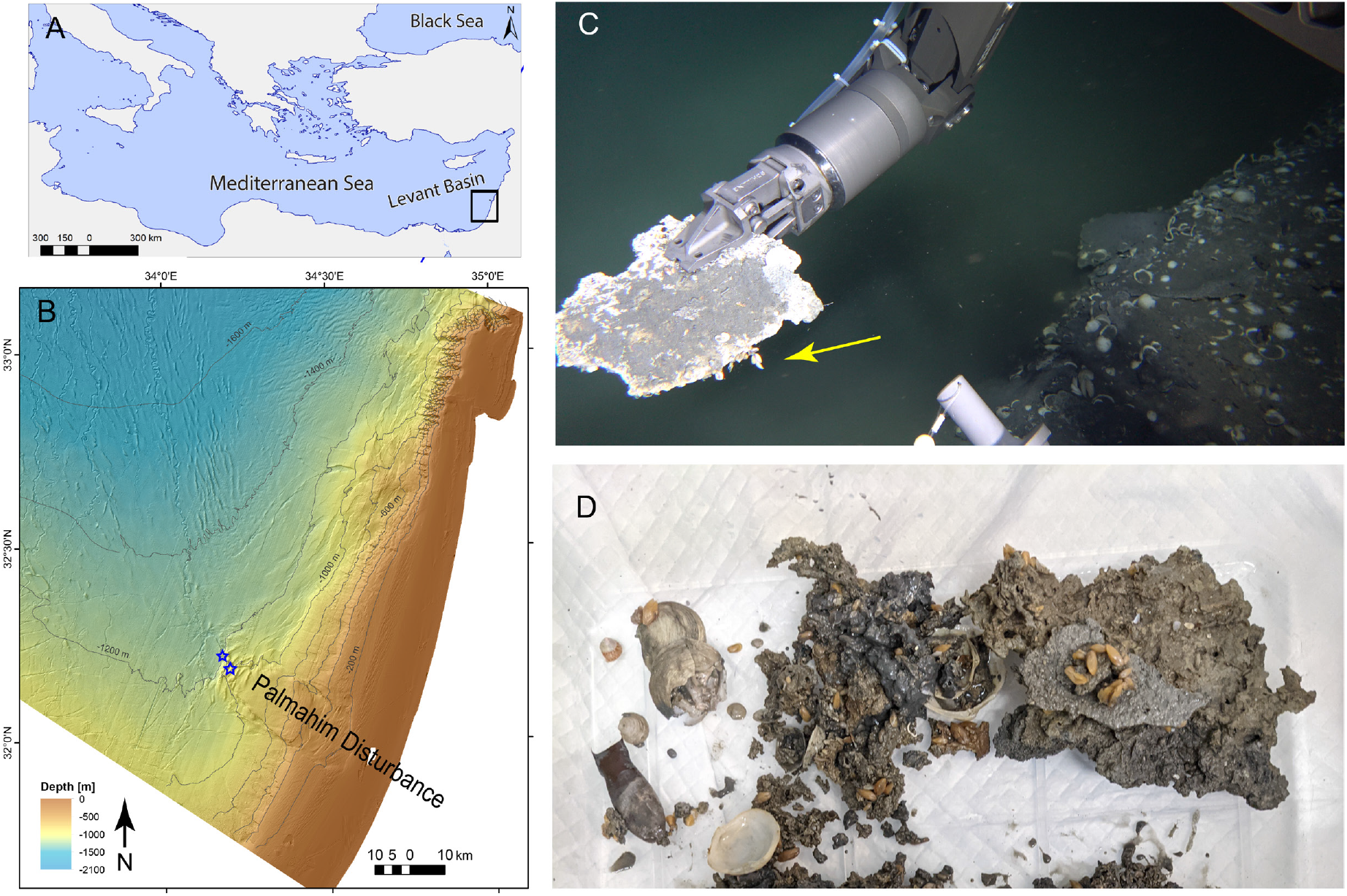
Collection sites of *Idas* specimens in the Eastern Mediterranean Palmahim Disturbance site (A,B, blue stars mark the two collection sites). *Idas* individuals were often found attached to authigenic carbonates (C-in situ, D - onboard).

**Figure S2:**
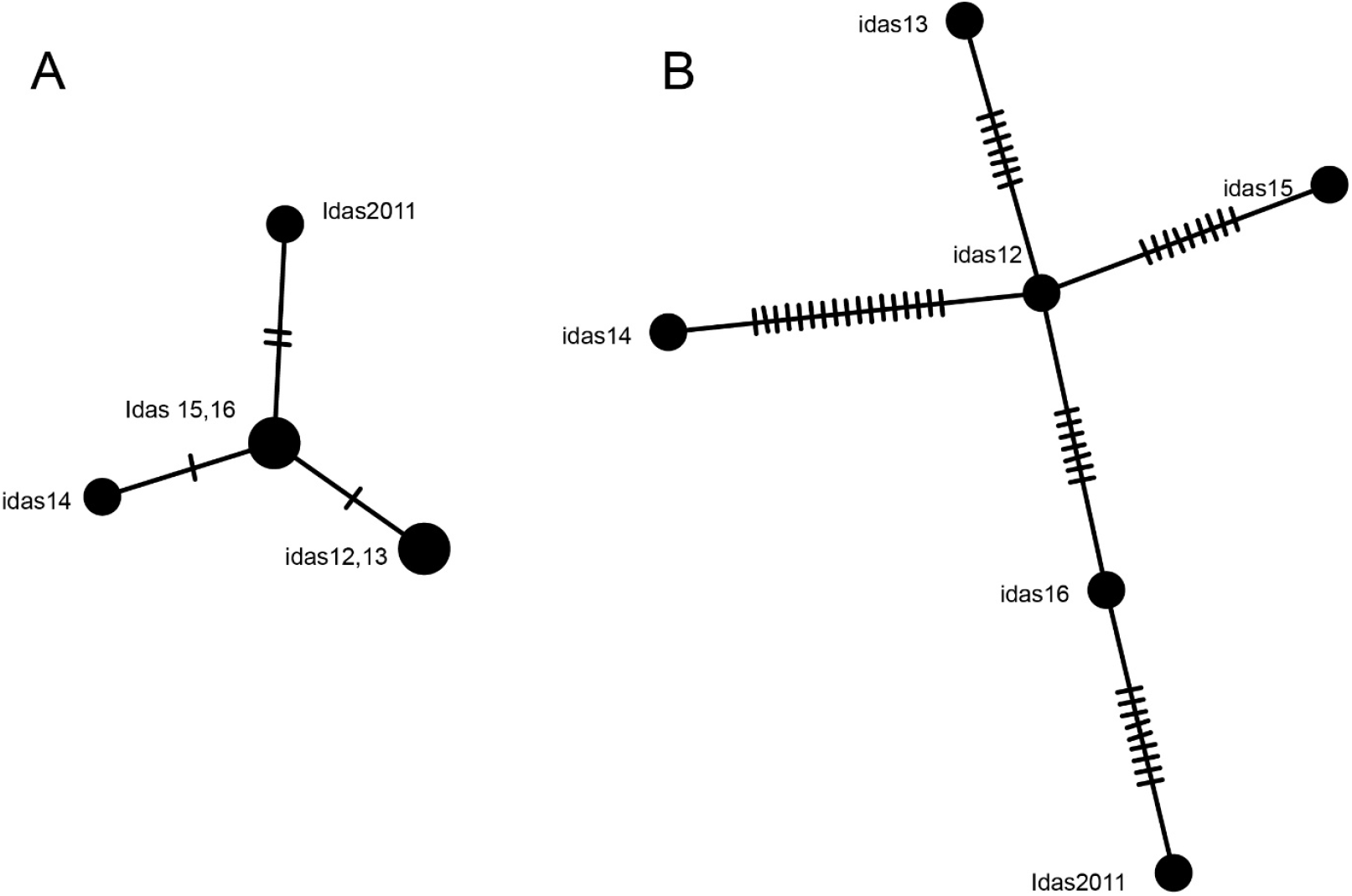
Mitochondrially encoded cytochrome c oxidase I (A) and mitochondrial genome (B) - based haplotype distances of the six *Idas modiolaeformis* individuals for which metagenomes were seqeuenced in this study. Each line indicates one single nucleotide polymorphism. The figure was produced with POPART (https://doi.org/10.1111/2041-210X.12410).

**Figure S3:**
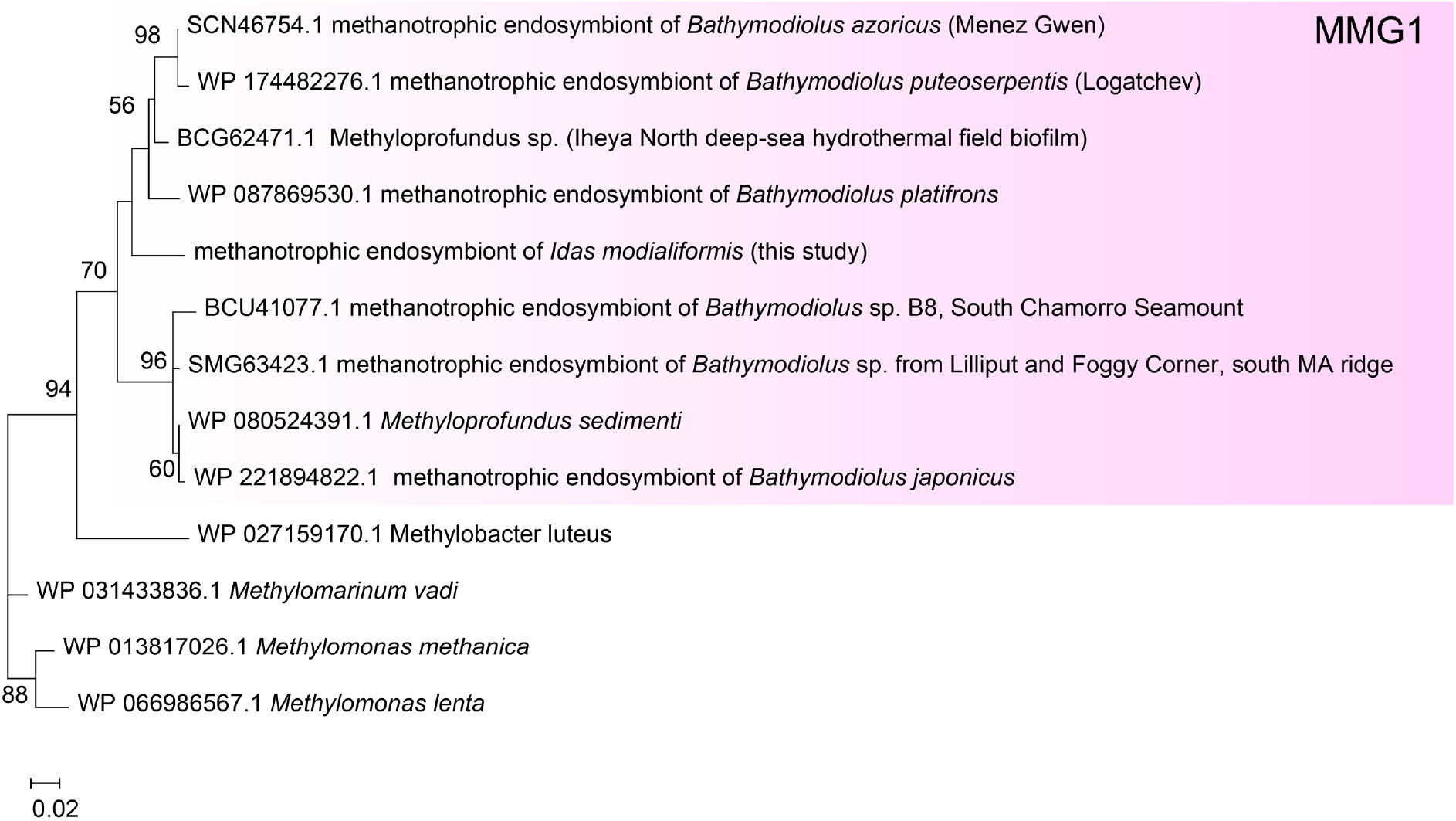
Phylogeny of the PmoA subunit of the particular methane monooxygenase of methane oxidizers including the methane-oxidizing symbiont of *Idas modiolaeformis*. The maximum likelihood midpoint-rooted tree is based on the LG model (MEGA11). The tree scale represents the number of substitutions per site.

**Figure S4:**
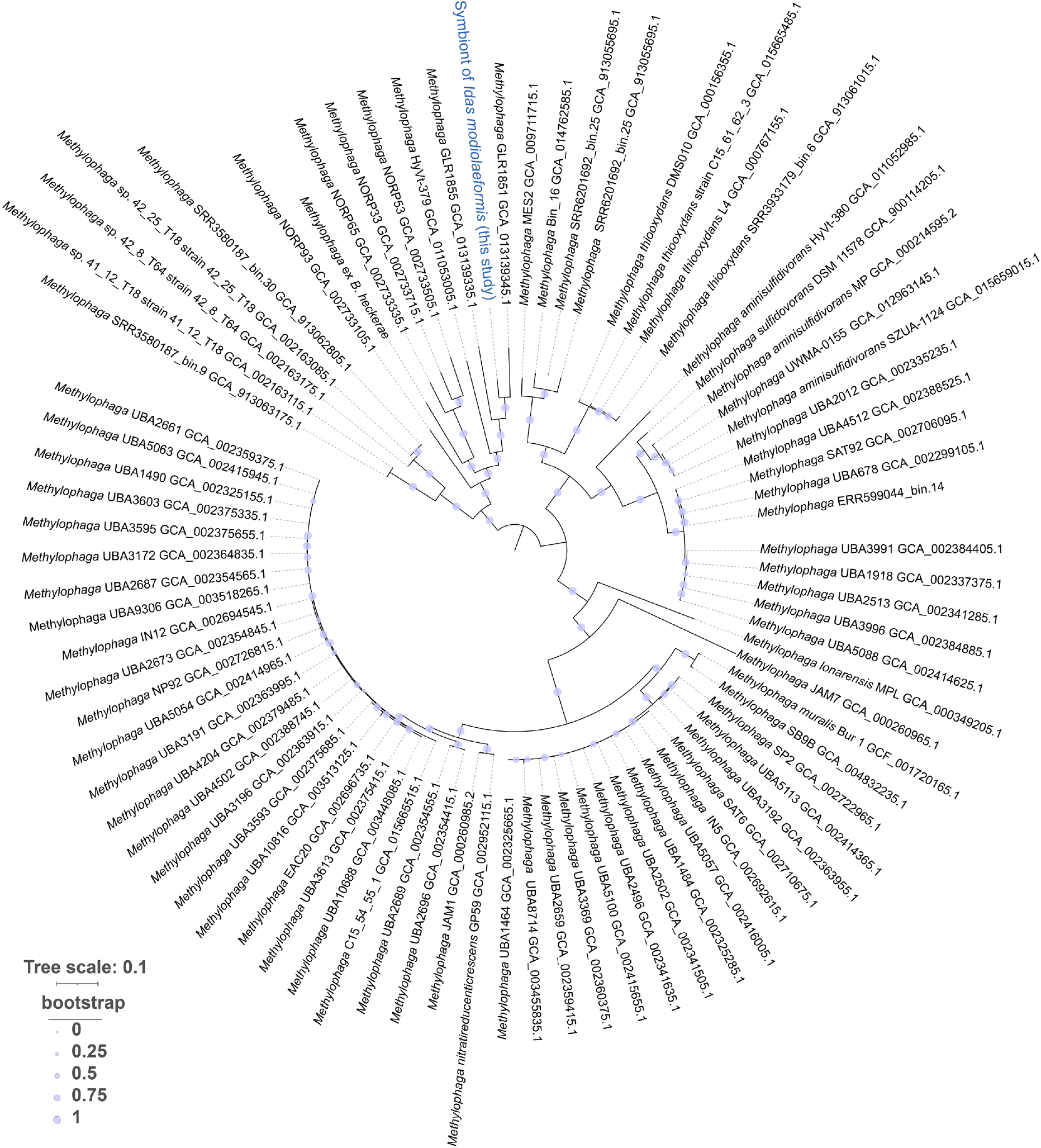
Phylogenomic tree of Methylophagaceae metagenome-assembled genomes, using the alignment of 172 protein sequences common to most gammaproteobacteria. The FastTree maximum likelihood tree was inferred using the JTT model, CAT approximation with 20 rate categories.

**Figure S5:**
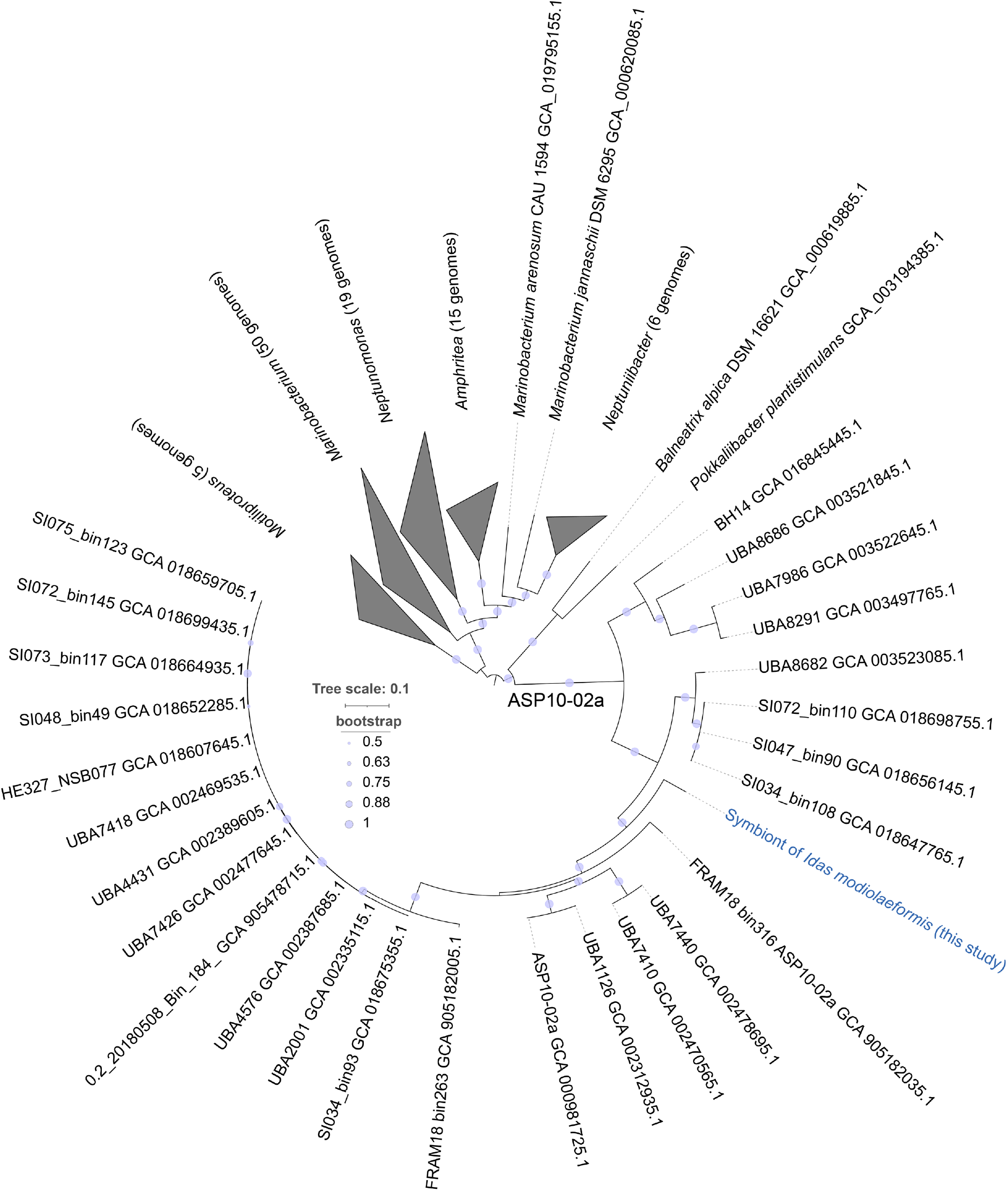
Phylogenomic tree of Nitrincolaceae metagenome-assembled genome, using the alignment of 172 protein sequences common to gammaproteobacteria. The FastTree maximum likelihood tree was inferred using the JTT model, CAT approximation with 20 rate categories.

**Figure S6:**
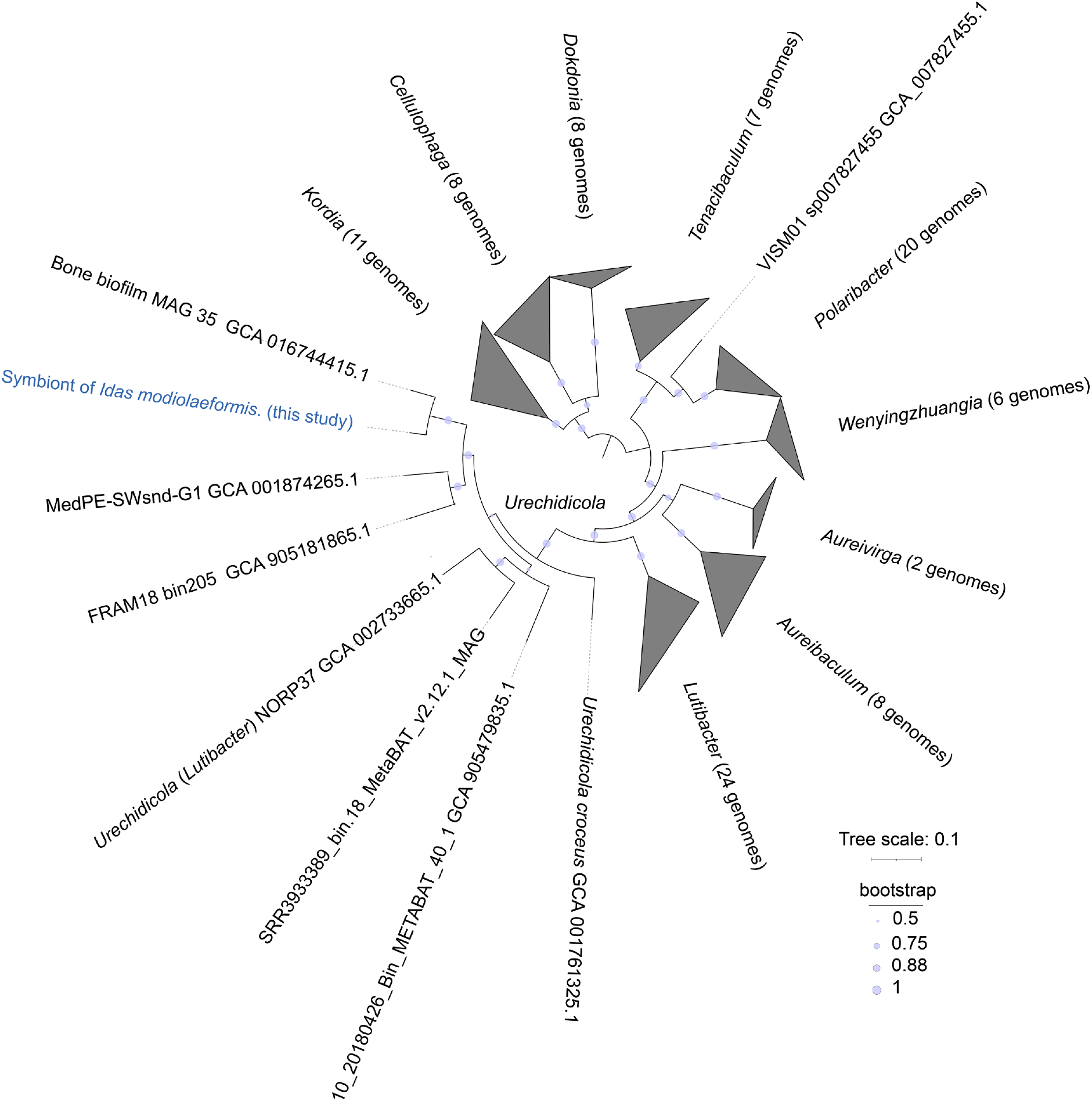
Phylogenomic tree of Flavobacteriaceae metagenome-assembled genome, using the alignment of 74 protein sequences common to bacteria. The FastTree maximum likelihood tree was inferred using the JTT model, CAT approximation with 20 rate categories.

**Figure S7:**
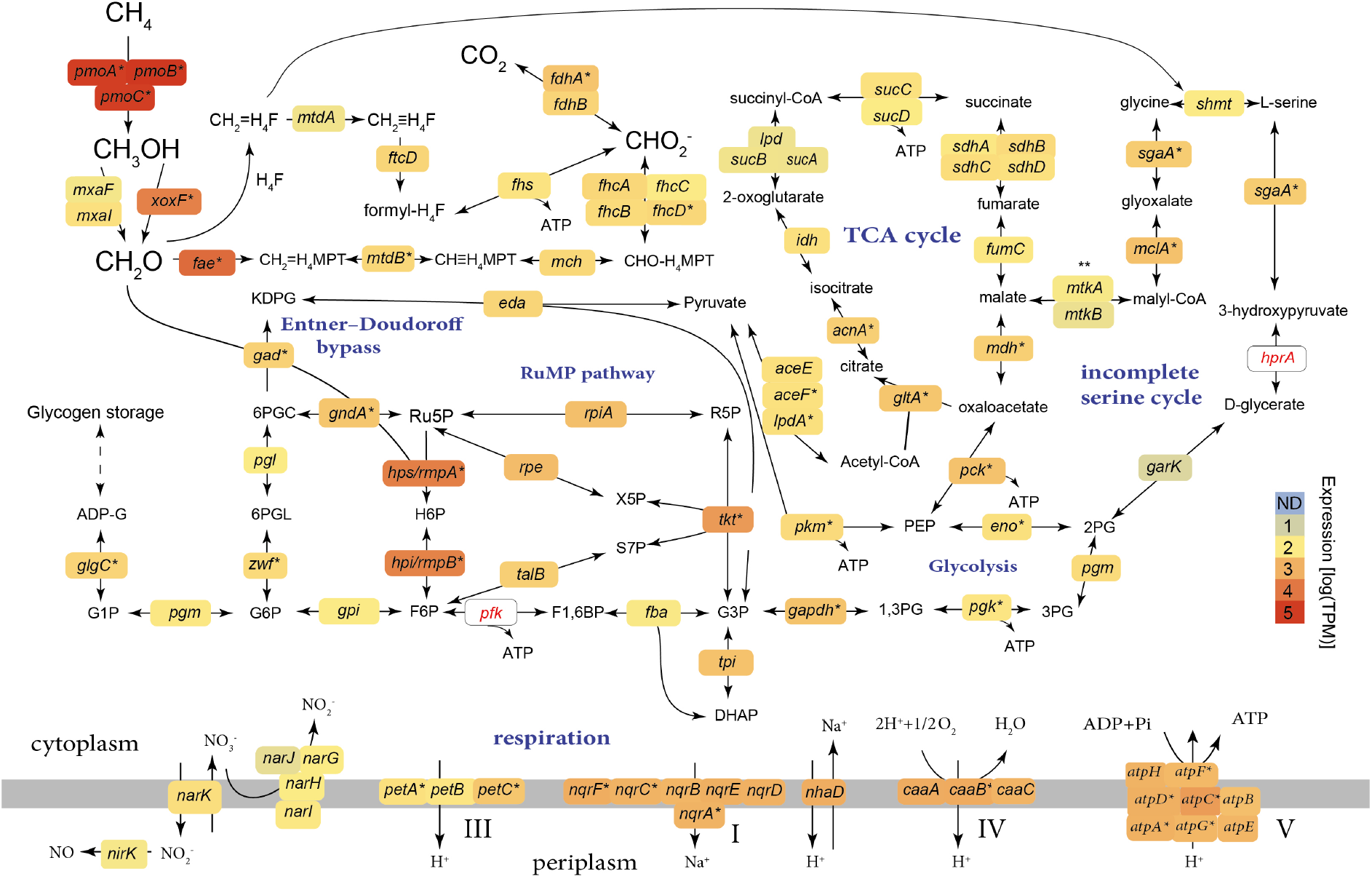
Central carbon metabolism in *Methyloprofundus* symbiont of *Idas modiolaeformis* and measured transcription levels in the metatranscriptomes. The *pfk* and *hprA* genes were not found in the MAG, therefore the Embden–Meyerhof–Parnas (EMP) variant of the ribulose monophosphate (RuMP) pathway is likely not present and the serine cycle is incomplete. The genes are as follows: calciumdependant methanol dehydrogenase *mxaFI*; lanthandine-dependant methanol dehydrogenase *xoxF*; 3-hexulose-6-phosphate synthase *hps/rmpA*; 3-hexulose-6-phosphate isomerase *hpi/rmpB*; transketolase *tkt*; ribose-5-phosphate isomerase *rpiA*; phosphoribulokinase *prk*; ribulose-phosphate 3-epimerase *rpe*; transaldolase *talB*; ATP-dependent 6-phosphofructokinase *pfk*; fructose-1,6-bisphosphate aldolase/phosphatase *fbp*: glucose-6-phosphate isomerase *gpi*; glucose-6-phosphate 1-dehydrogenase *zwf*; 6-phosphogluconolactonase *pgl*; phosphogluconate dehydratase *edd*; 2-dehydro-3-deoxy-phosphogluconate/2-dehydro-3-deoxy-6-phosphogalactonate aldolase *eda*; phosphoglucomutase *pgm*; glucose-1-phosphate adenylyltransferase *glgC*; fuctose-bisphosphate aldolase *fba*; triosephosphate isomerase *tpi*; formaldehyde activating enzyme *fae*; methylene tetrahydromethanopterin dehydrogenase *mtdB*; methenyltetrahydromethanopterin cyclohydrolase *mch*; formyltransferase/hydrolase complex *fhcABCD*; NAD(P)-dependent methylenetetrahydromethanopterin dehydrogenase *mtdA*; bifunctional methylenetetrahydrofolate dehydrogenase / methenyltetrahydrofolate cyclohydrolase *fold*; formate--tetra hydrofol ate ligase *fhs*; formate dehydrogenase *fdhAB*; glyceraldehyde-3-phosphate dehydrogenase *gapdh*; phosphoglycerate kinase *pgk*; phosphoglucomutase *pgm*; enolase *eno*; phosphoenolpyruvate synthase *ppsA*; phosphoenolpyruvate carboxykinase *pck*; pyruvate dehydrogenase *aceEF-lpdA*; pyruvate kinase *pkm*; citrate synthase *gltA*; aconitase *acnA*; isocitrate dehydrogenase *idh*; 2-oxoglutarate dehydrogenase complex *sucAB-lpd*; succinate--CoA ligase *sucCD*; succinate dehydrogenase *sdhABC*; fumarate hydratase class II *fumC*; malate hydrogenase *mdh*; malate--CoA ligase *mtkAB*; malyl-CoA lyase *mclA*; serine--glyoxylate aminotransferase *sgaA*; serine hydroxymethyltransferase *shmt*; glycerate dehydrogenase *hprA*; glycerate 2-kinase *garK*; caa_3_-type cytochrome c oxidase *cyoABC*; ubiquinol-cytochrome c reductase *petABC*; Na(+)-translocating NADH-quinone reductase *nqrA-F*; Na(+)/H(+) antiporter *nhaD*; respiratory nitrate reductase *narGHIJ*, copper-containing nitrite reductase *nirK*; nitrate/nitrite antiporter *narK*; ATP synthase *atpA-F*. Metabolites: OA, oxaloacetate; PEP, phosphoenolpyruvate; 2-phosphoglycerate, 2PG; 3-phosphoglycerate, 3PG; 1,3-bisphosphoglycerate 1,3BPG; 3-phosphoglyceraldehyde, G3P; dihydroxyacetone phosphate, DHAP; fructose 1,6-bisphosphate, F1,6BP; fuctose 6-phosphate, F6P; hexulose 6-phosphate, H6P; ribulose 5-phosphate, Ru5P; ribulose-1,5-bisphosphate, Ru1,5BP; ribose 5-phosphate, R5P; glucose 6-phosphate, G6P; 6-phosphogluconolactonase, 6PGL; 2-Dehydro-3-deoxy-D-gluconate 6-phosphate, 6PGC; 2-keto-3-deoxy-6-phosphogluconate, KPDG; D-Xylulose 5-phosphate, X5P; sedoheptulose 7-phosphate, S7P; gucose 1-Phosphate, G1P; ADP-glucose, ADP-G; tetrahydrofolate, H4F; tetrahydromethanopterin, H4MPT. Average expression values from 4 individuals are shown.

**Figure S8:**
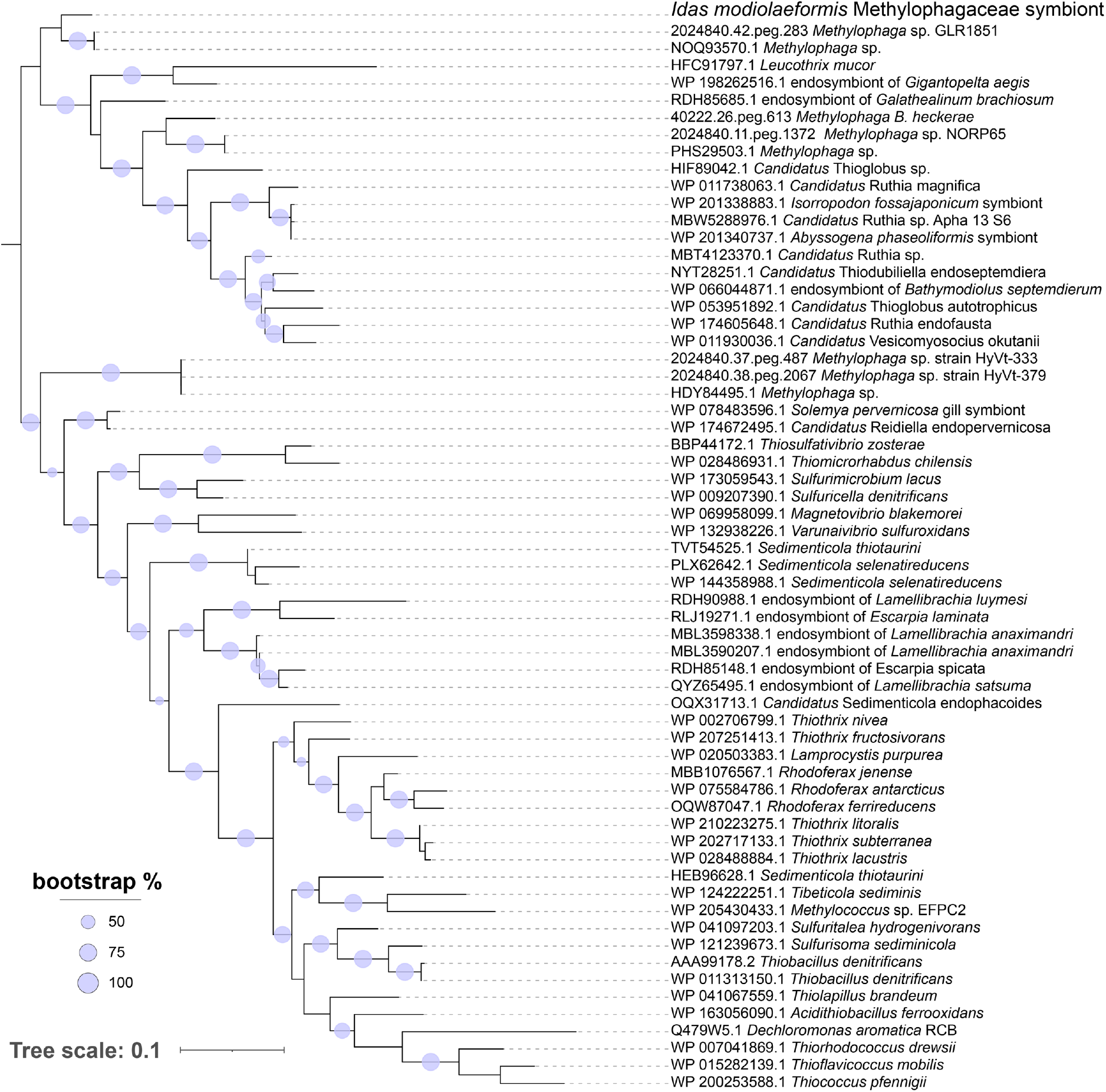
Phylogenetic tree of the CbbM (form II Ribulose-1,5-bisphosphate carboxylase-oxygenase, RuBisCO) amino acid sequences from symbiotic Methylophagaceae and selected bacteria. The maximum likelihood tree is based on the LG+I+G4 model (IQ-TREE 2). The tree scale represents the number of substitutions per site.

**Figure S9:**
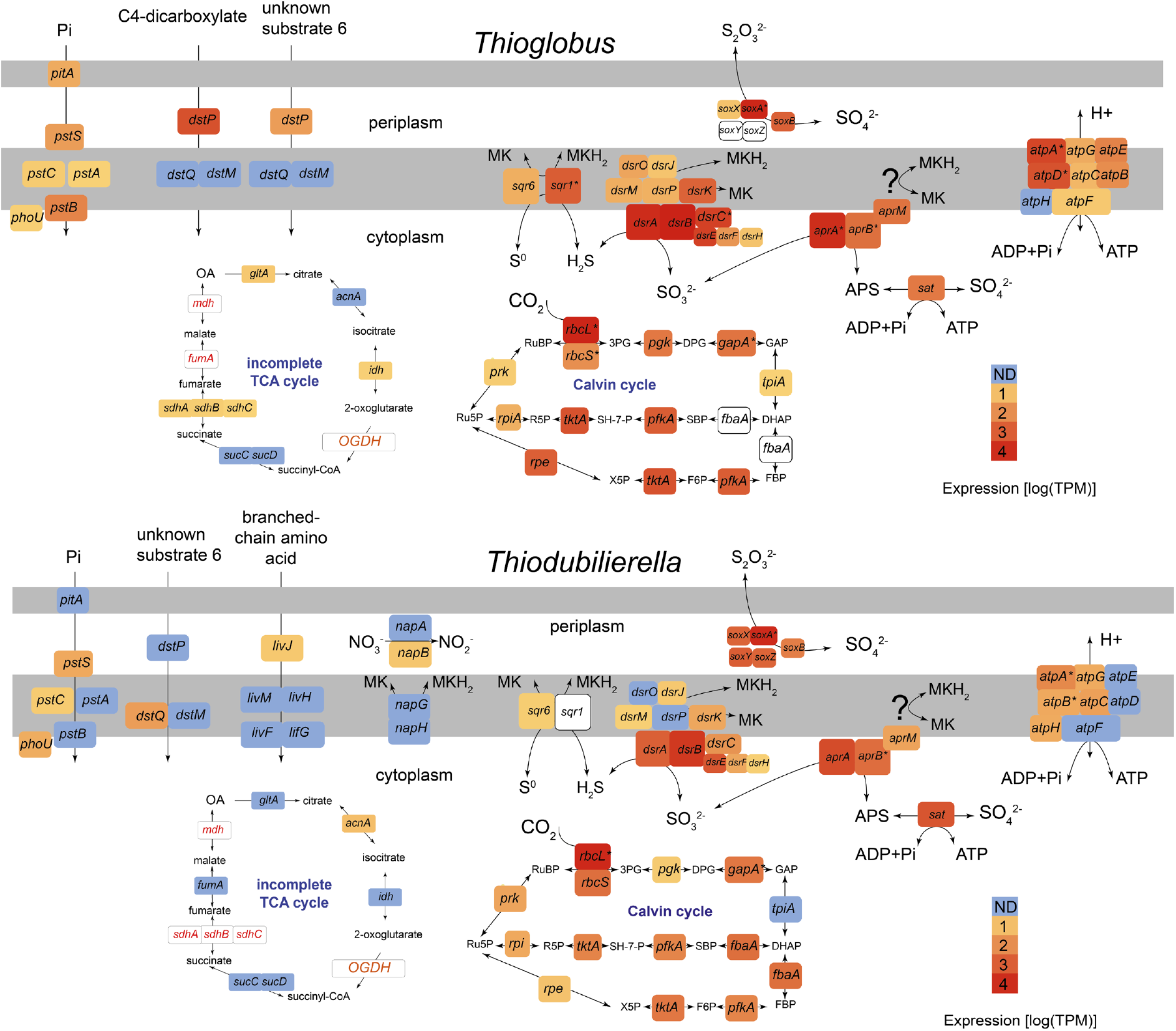
Central metabolism in the sulfur-oxidizing symbionts of *Idas modiolaeformis* (top panel: *Thioglobus*, bottom panel: *Thiodubilierella*). White boxes indicate functions that were not found. Expression levels shown correspond to transcription levels from the metatranscriptomic data. Average expression values from 4 individuals are shown. Proteins/genes: sulfur oxidation complex *soxAXYZB*; sulfide:quinone oxidoreductase, type I/VI, *sqr1* / 6; dissimilatory sulfite reductase *dsrAB*; sulfite reduction-associated complex *dsrMKJOP-C-EFH*; adenylylsulfate reductase subunit *aprABM*; ATP synthase *atpA-F*; ribulose bisphosphate carboxylase form I *rbcLS*; phosphoglycerate kinase *pgk*; glyceraldehyde-3-phosphate dehydrogenase *gapA*; triosephosphate isomerase *tpiA*; ribose-5-phosphate isomerase *rpi*; tranketolase *tktA*; pyrophosphate-dependent fructose 6-phosphate-1-kinase *pfkA*; ribulose-phosphate 3-epimerase *rpe*; fructose-bisphosphate aldolase class *fbaA*; citrate synthase *gltA*; aconitase *acnA*; isocitrate dehydrogenase *idh*; 2-oxoglutarate dehydrogenase complex *OGDH*; succinate--CoA ligase *sucCD*; succinate dehydrogenase *sdhABC*; fumarate hydratase class I *fumA*; malate hydrogenase *mdh*. Metabolites: OA, oxaloacetate; 3-Phosphoglyceric acid, 3PG, glyceraldehyde 3-phosphate, GAP; 1,3-bisphosphoglycerate, DPG; dihydroxyacetone phosphate, DHAP; fructose 1,6-bisphosphate, FBP; fuctose 6-phosphate, F6P; D-Xylulose 5-phosphate, X5P; sedoheptulose-1,7-bisphosphate, SBP; sedoheptulose 7-phosphate, SH-7-P; ribose 5-phosphate, R5P; Ru5P; ribulose-1,5-bisphosphate, RuBP. MK/MKH_2_ represents the oxidized and reduced quinone pool. Pi is pyrophosphate.

**Table S1 (Excel sheets)**: Genomic features of *Idas* symbionts, and their RNA- and protein-level expression (raw mapped read counts and log[transcripts per million x 10^6^] for RNA, % normalized spectral abundance factor, %NSAF for proteins). Each one of the six metagenome-assembled genomes is summarized in a separate tab. Coding sequences, as well as their protein-level translations, are included. Rast SEED annotations are shown.

## Notes

### Competing Interest Statement

The authors have declared no competing interest.

